# Miniaturized subcutaneous cellular implants for sustained therapeutic protein delivery in resource-limited settings

**DOI:** 10.64898/2026.06.03.730028

**Authors:** Minseok Lee, Bo Wang, Kecheng Wang, Kento Okada, James Arthur Flanders, Alexander Barutis, Juan M. Melero-Martin, Minglin Ma

## Abstract

Cell encapsulation offers a promising strategy for sustained therapeutic protein delivery, obviating the need for repeated injections. Among potential implantation sites, the subcutaneous space is particularly attractive for its accessibility and amenability to minimally invasive procedures. However, performance of subcutaneous devices reported to date has been limited due to various challenges including foreign body response (FBR) and inadequate mass transfer. Moreover, typical encapsulation devices require surgeries for implantation and retrieval, limiting their potential use in resource-limited settings. Here we present a miniaturized cell encapsulation platform comprising cells engineered to produce therapeutic proteins and an FBR-mitigating zwitterionic polyurethane nanofibrous membrane, in a thin cylindrical form factor compatible with applicator-based minimally invasive implantation and retrieval. Clonal mesenchymal stromal cells engineered to produce PGT121, a broadly neutralizing anti-HIV-1 antibody, were encapsulated and inserted subcutaneously, achieving long-term cell survival and sustained serum PGT121 concentrations for up to 36 weeks across multiple murine models. Cell-loaded devices retained therapeutic function after cryopreservation, supporting their potential use as an off-the-shelf product that can be centrally manufactured and implanted on-site without specialized infrastructure. The custom-designed applicator-based implantation and minimally invasive retrieval procedures were demonstrated in a more clinically relevant minipig model. These mini-“cellular factories” represent a translatable strategy for sustained delivery of biologic drugs in resource-limited settings.

**One Sentence Summary:** An insertable and retrievable mini cellular construct enables sustained protein delivery, supporting its potential use in resource-limited settings.

## INTRODUCTION

Biologic drugs, including monoclonal antibodies, hormones, and other recombinant proteins, have become a dominant class of therapeutics over the past two decades, now accounting for approximately one-third of all US Food and Drug Administration (FDA)-approved new drugs annually(*1, 2*). However, most of them require repeated injections owing to their limited circulating half-lives or high required doses(*3*). Despite decades of engineering efforts to extend dosing intervals, the fundamental need for chronic repeated administration persists, motivating the development of cell-based platforms that function as implantable protein factories capable of sustained therapeutic protein secretion *in vivo*(*4*).

Advances in biological engineering, including genome and epigenome editing, have expanded cell-based therapeutic platforms beyond the native secretory functions of implanted cells, enabling genetically engineered cell-based implants as a versatile approach to sustained biologic delivery(*5, 6*). However, a central challenge for cell-based therapies is host immune rejection of the implanted cells, particularly when using allogeneic cell sources, which are strongly preferred over autologous cells for their off-the-shelf availability, scalable manufacturing, and lower per-patient cost(*6*). Systemic immunosuppression can mitigate rejection but carries significant side effects, including increased risk of infection and malignancy. The cell encapsulation approach addresses these limitations by confining allogeneic therapeutic cells within semi-permeable membranes, which provide immunoprotection without systemic immunosuppression, permit facile mass transfer of nutrients, oxygen, and secreted therapeutic proteins, and offer a level of safety and controllability through physical containment and device retrievability. The recent FDA approval of Encelto, the first and only approved genetically engineered cell encapsulation product, validates the clinical feasibility of this platform as a therapeutic approach(*7*). This clinical success, however, was enabled by a uniquely permissive therapeutic context. Intravitreal implantation places the device within an immune-privileged, anatomically confined compartment that sustains local protein concentrations within the therapeutic range at the site of action, rendering nanogram-per-day protein secretion from the implant sufficient for therapeutic efficacy(*8, 9*). Extending this platform to broader therapeutic applications, particularly those requiring systemic protein delivery from non-immune-privileged sites, remains a significant unmet need. A primary challenge is the FBR and fibrotic reactions that lead to a dense fibrotic layer impeding mass transfer of nutrients, oxygen, and therapeutic proteins. This fibrotic reaction has limited long-term device function in multiple clinical cell encapsulation trials, particularly those involving islet encapsulation(*10, 11*). A second challenge is the procedural complexity associated with surgical implantation. This challenge is particularly acute in resource-limited settings, where access to surgical infrastructure and trained personnel is often limited. The subcutaneous space has emerged as a promising implantation site that addresses this procedural challenge, owing to its accessibility for minimally invasive procedures(*12*). Nevertheless, the subcutaneous microenvironment is inherently limited by low oxygen availability, low vascular density, and reactive local immune responses, all of which compromise long-term device function(*13*).

Here, we present a miniaturized cellular implant for long-term therapeutic protein delivery via subcutaneous implantation, with a focus on accessibility in resource-limited settings. The thin, cylindrical device form factor and implantation procedure were inspired by subcutaneous contraceptive implants, which have been widely adopted in resource-limited settings owing to their simplicity of insertion and retrieval(*14*). The device features an electrospun nanofibrous membrane of zwitterionically modified polyurethane (ZPU), providing immunoprotection, mechanical robustness and FBR-mitigating properties. As a model for therapeutic cells, we engineered mesenchymal stromal cells (MSCs) to secrete PGT121, a broadly neutralizing antibody (bNAb) under clinical evaluation for HIV-1 prevention and treatment(*15*). We demonstrated sustained cell survival and PGT121 production for up to 36 weeks in the subcutaneous space across multiple mouse models, with retrieved devices maintaining structural integrity, immunoprotective function, and minimal fibrotic reactions. The platform supported off-the-shelf use through cryopreservation and extended to human induced pluripotent stem cell-derived MSCs (hiMSCs) as a clinically relevant cell source. Applicator-based implantation and minimally invasive retrieval were further validated at clinically relevant scale in a minipig model. Collectively, these results establish a translatable strategy for sustained biologic delivery via cellular implants in resource-limited clinical settings.

## RESULTS

### Design and optimization of zwitterionically modified polyurethane (ZPU) nanofibrous encapsulation device

Our workflow for therapeutic cell delivery (Fig. 1A) comprises device fabrication, engineering of therapeutic protein-producing cells, cell loading into the device, and subcutaneous implantation. We designed the cell encapsulation device to meet five key design criteria (Fig. 1B): (i) immunoprotection and cell confinement through an electrospun nanofibrous membrane, (ii) FBR mitigation and facile mass transfer through zwitterionic surface chemistry, (iii) mechanical robustness sufficient to preserve structural integrity after implantation, (iv) a cylindrical geometry compatible with a custom-designed applicator for minimally invasive implantation, and (v) cryopreservability for off-the-shelf use.

**Fig. 1.**
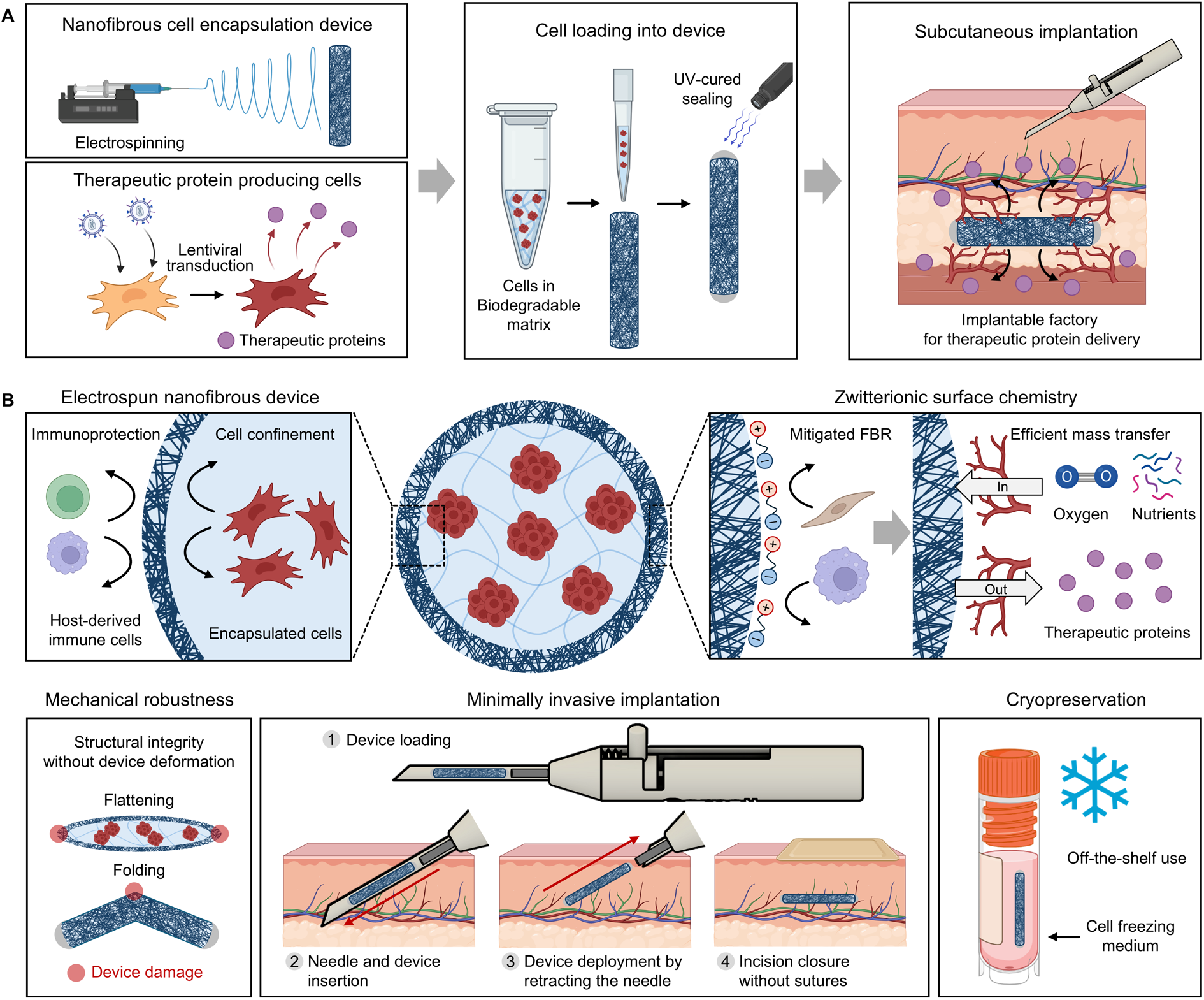
Design and workflow of the ZPU cell encapsulation platform. **(A)** Schematic of the therapeutic cell delivery workflow. **(B)** Schematic of the ZPU cell encapsulation device illustrating its five key design criteria.

Building on the low FBR and facile mass transfer previously achieved with ZPU in islet encapsulation(*16*), we aimed to expand this approach to genetically engineered therapeutic protein-producing cells for a broader range of therapeutic targets, while optimizing the device for subcutaneous space. ZPU was synthesized as previously described (Fig. S1A). The chemical structures of the synthesized SB-Diol (Fig. S1B), ZPU, and PU, a control synthesized without SB-Diol (Fig. S1C), were confirmed by ^1^H NMR spectroscopy. Cylindrical membranes were then fabricated by electrospinning ZPU or PU polymer solutions dissolved in 1,1,1,3,3,3-hexafluoro-2-propanol (HFIP). FT-IR analysis of the resulting membranes showed a strong peak at 1,105 cm^−1^ in both control PU and ZPU spectra, assigned to the C-O-C asymmetric stretching of the PTMG soft segment, whereas a characteristic peak at 1,037 cm^−1^, attributed to SO_3_^−^ symmetric stretching, was observed only in the ZPU spectrum, confirming the successful incorporation of sulfobetaine groups (Fig. 2A). The fiber diameter of the electrospun membrane was tunable by varying the concentration of the ZPU polymer solution, with higher concentrations yielding larger fiber diameters (Fig. 2B, Fig. S2A). We selected the membrane fabricated from 10% ZPU solution, which produced an average fiber diameter of 0.192 µm, as this fiber size was shown in our previous work to provide effective immunoprotection and facile mass transfer (Fig. 2C)(*16, 17*). Qualitatively, the cylindrical membrane exhibited mechanical elasticity, stretching to more than twice its original length (Fig. 2D). Tensile testing revealed an ultimate tensile strength of approximately 8 MPa and an elongation at break exceeding 200% (Fig. S2B). The device dimensions are readily tunable, with the diameter of the cylindrical membrane determined by the mandrel collector diameter during electrospinning and the length adjusted by cutting after collection (Fig. S2C). In this study, we selected a device with an inner diameter of 1.2 mm and a length of 2.5 cm. These dimensions are smaller than those of aforementioned clinically approved subcutaneous contraceptive implant Nexplanon (2 mm in diameter and 4 cm in length), preserving the advantages of minimally invasive implantation and retrieval procedures. The device is readily scalable to larger dimensions for higher therapeutic doses or larger species. To validate the immunoprotective function of the device and its ability to mitigate the material-driven FBR *in vivo*, acellular ZPU devices were implanted into the dorsal subcutaneous space of C57BL/6 mice. Throughout the 16 weeks implantation period, histological analysis confirmed no evidence of immune cell infiltration and minimal FBR, with the fibrotic layer around the device consisting of only a few cell layers (Fig. 2E).

**Fig. 2.**
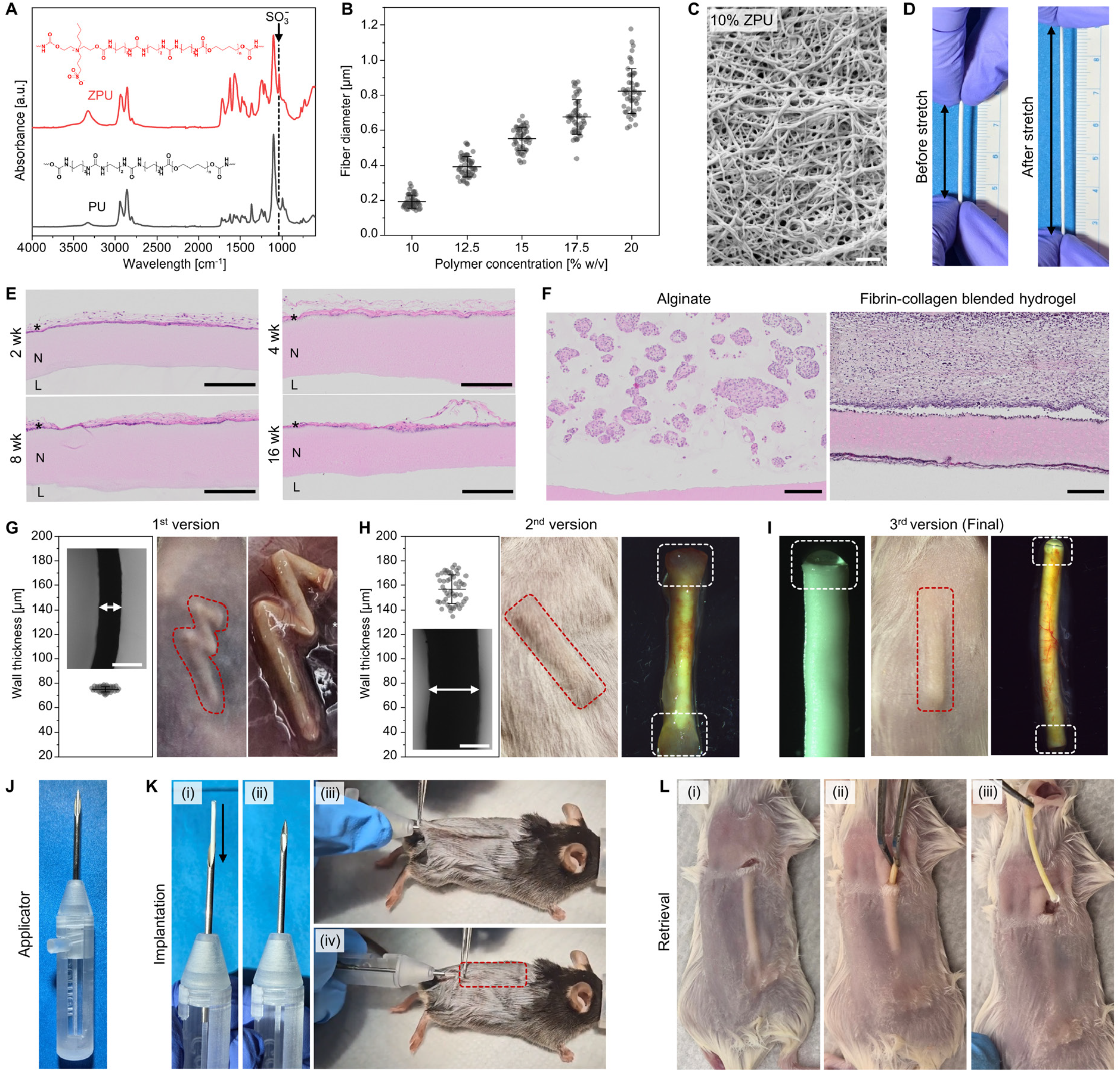
Design and optimization of the ZPU nanofibrous encapsulation device. **(A)** FT-IR spectra of ZPU (red) and PU (black) membranes. The dashed line indicates the SO_3_^−^ stretching peak at 1,037 cm^−1^. **(B)** Fiber diameter of ZPU membranes fabricated from ZPU solutions at various concentrations (10, 12.5, 15, 17.5, and 20% w/v in HFIP). (*n = 50*) **(C)** Representative SEM image of an electrospun ZPU membrane (10% w/v ZPU solution). Scale bar, 2 µm. **(D)** Photographs of the ZPU device before (left) and after (right) stretching. **(E)** H&E-stained cross-sectional images of acellular ZPU devices retrieved at 2, 4, 8, and 16 weeks post-implantation in C57BL/6 mice. L, device lumen; N, nanofibrous membrane; *, fibrotic layer. Scale bar, 200 µm. (*n = 3*) **(F)** H&E-stained cross-sectional images of devices encapsulating cells mixed with alginate (left; retrieved at 93 days) (*n = 5*) or a fibrin-collagen blended hydrogel (right; retrieved at 120 days) (*n = 3*) from RAG2-KO mice. Scale bar, 200 µm. **(G–I)** Iterative device optimization (Ver 1–3). **(G)** Ver 1: wall cross-section (left; scale bar, 100 µm) and thickness quantification (*n = 50*), implantation site (middle), and retrieved device at 8 weeks (right) (*n = 5*). **(H)** Ver 2: wall cross-section (left; scale bar, 100 µm) and thickness quantification (*n = 50*), implantation site (middle), and retrieved device at 4 weeks (right) (*n = 5*). **(I)** Ver 3: end cap (left), implantation site (middle), and retrieved device at 16 weeks (right) (*n = 3*). All devices implanted subcutaneously in BALB/c mice. **(J)** Photograph of the 3D-printed applicator. (**K**) Implantation procedure: (i-ii) device loading, (iii) needle insertion, and (iv) deployment. **(L)** Retrieval procedure: (i) skin incision, (ii) partial dissection and grasping, and (iii) extraction.

Three key device parameters were further optimized for genetically engineered therapeutic cells in the subcutaneous space: the matrix core, wall thickness, and sealing method. In our previous works, a non-degradable alginate matrix was used to maintain uniform distribution of non-proliferative islets. Here, we replaced it with a biodegradable fibrin-collagen blended hydrogel, which provides cell-adhesive cues and allows proliferative cells to expand within the device as the matrix degrades, maximizing therapeutic protein productivity(*18*). H&E-stained cross-sectional images of devices retrieved from mice confirmed this rationale: MSC spheroids in alginate retained their original morphology (Fig. 2F, left), whereas those in the degradable matrix expanded throughout the device lumen (Fig. 2F, right). We then tested three device iterations to determine the optimal wall thickness and sealing method. The first version, with a wall thickness of ∼75 µm and thermally sealed ends, exhibited deformation, posing a risk of structural defects (Fig. 2G, S2D). In the second version, wall thickness was increased to ∼ 157 µm to improve mechanical robustness. While this device showed improved mechanical stability, the thermal sealing still resulted in sharp and rough edges that promoted fibrotic reactions and occasionally introduced structural defects at the sealed ends due to uneven thinning (Fig. 2H, S2E). In the final version, we replaced thermal sealing with a photocurable adhesive, which yielded smooth and uniform end caps. This device maintained mechanical robustness with no evidence of deformation or sealing defects over 16 weeks post-implantation (Fig. 2I). Inspired by applicators used for subcutaneous contraceptive implants, we then designed a 3D-printed custom-designed applicator comprising a body with a locking notch, a needle with slider, and an obturator (Fig. 2J), which together with the cylindrical form factor enabled minimally invasive subcutaneous implantation (Fig. 2K) and retrieval (Fig. 2L) without specialized surgical training.

### Engineering of PGT121-producing MSCs and *in vitro* device evaluation

Fig. 3A illustrates the genetic engineering workflow for producing therapeutic protein-secreting MSCs, including transfection, transduction, single cell cloning, and high producing clone selection. We selected MSCs as the therapeutic cell type given their established clinical safety profile, low immunogenicity, and immunomodulatory properties, which together support their use in allogeneic applications (*19, 20*). We engineered bone marrow-derived MSCs from C57BL/6 mice to constitutively secrete PGT121 by lentiviral transduction, which enables stable transgene integration (Fig. S3A). Stably transduced cells were enriched by antibiotic selection, yielding a polyclonal population that produced PGT121 at 4.79 µg per 10^6^ cells per day (Fig. 3B). Flow cytometric analysis confirmed that both control and engineered MSCs retained characteristic MSC surface marker profiles (Fig. S3B, S3C). We then performed single-cell cloning to isolate clones with high protein production and reduced functional heterogeneity. Transduced MSCs were sorted into 96-well plates (276 wells total) and expanded clonally, and PGT121 secretion was assessed by primary (Fig. S3D) and secondary ELISA screening (Fig. 3B). The highest-producing clone (clone #6, producing PGT121 at 30.04 µg per 10^6^ cells per day) was selected for subsequent studies. Cumulative population doubling level (CPDL) analysis over eight consecutive passages confirmed that clone #6 maintained proliferative capacity without progressive decline, indicating that neither lentiviral engineering nor clonal selection compromised proliferative capacity (Fig. 3C).

**Fig. 3.**
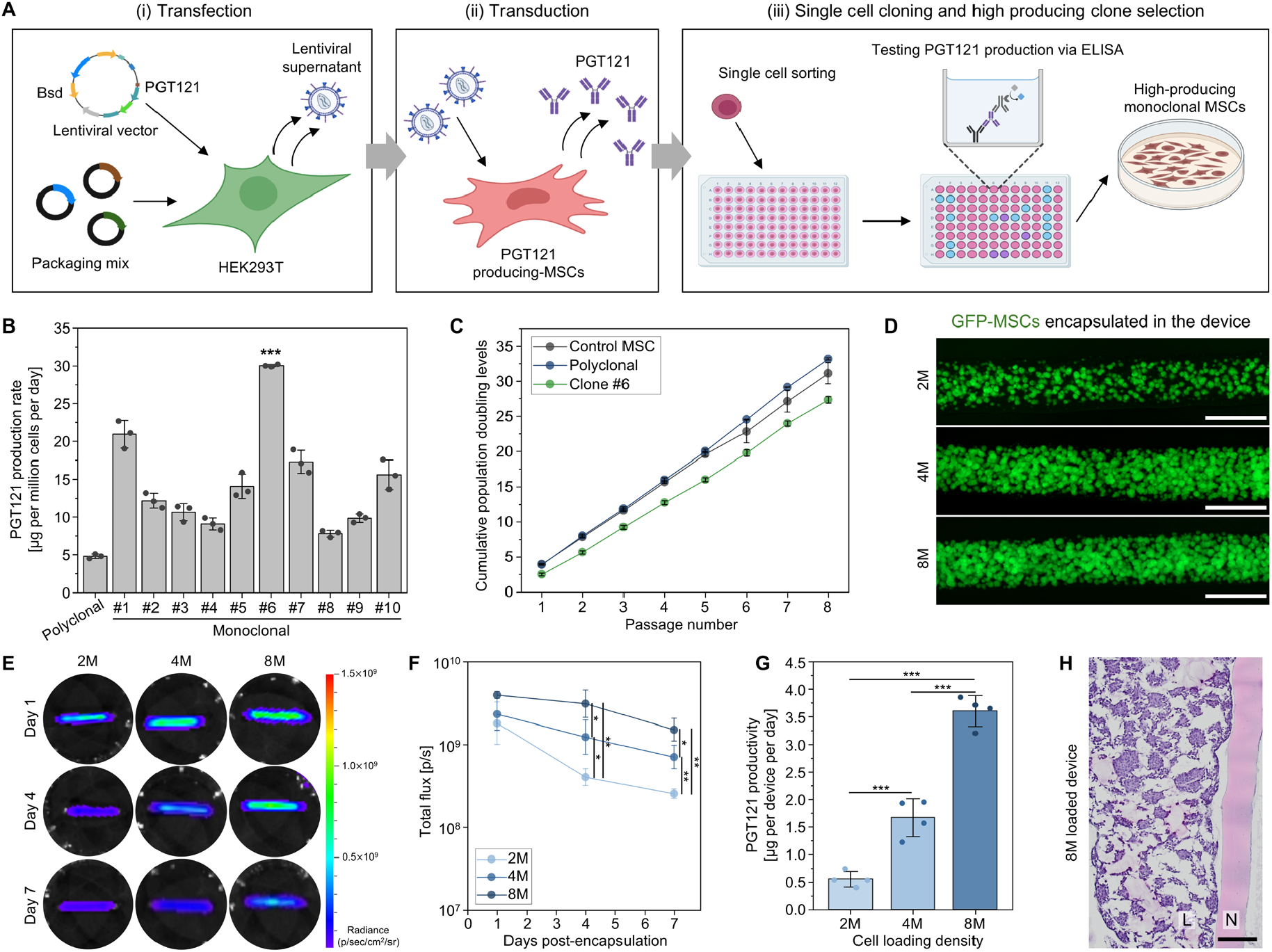
Engineering of PGT121-producing MSCs and *in vitro* device evaluation. **(A)** Schematic of the genetic engineering workflow for producing PGT121-secreting MSCs. **(B)** Per-cell PGT121 productivity of the engineered polyclonal MSCs and ten high-producing clones. *\*\*\*p* < 0.001, clone #6 compared to all other groups. **(C)** Cumulative population doubling levels over passage of control MSCs, polyclonal PGT121-producing MSCs, and clone #6 (*n = 3*). **(D)** Fluorescence images showing 2, 4, or 8 × 10^6^ GFP-MSC spheroids loaded into devices. Scale bar, 1,000 µm. **(E)** Bioluminescence images of devices loaded with 2, 4, or 8 × 10^6^ PGT121/GFP/Luc-MSCs and **(F)** corresponding quantification (*n = 4*) at days 1, 4, and 7 post-encapsulation. *\*p* < 0.05, *\*\*p* < 0.01. **(G)** *In vitro* per-device PGT121 productivity measured between days 4 and 5 post-encapsulation (*n = 4*). *\*\*\*p* < 0.001. **(H)** H&E-stained cross-sectional images of devices loaded with 8 × 10^6^ MSCs after 1 week of *in vitro* culture. L, device lumen; N, nanofibrous membrane. Scale bar, 200 µm.

Engineered MSCs were then encapsulated into the ZPU device and cultured *in vitro* to evaluate cell survival and PGT121 secretion after encapsulation. Prior to encapsulation, MSC spheroids were prepared to enhance cell survival through increased cell–cell and cell–ECM interactions(*21*). Immunofluorescence imaging of devices loaded with 2, 4, or 8 × 10^6^ GFP-MSCs, following removal of the device membrane to expose the lumen, confirmed uniform cell distribution (Fig. 3D). Cell survival was monitored by bioluminescence imaging (BLI), and *in vitro* PGT121 productivity per device was quantified by ELISA. Encapsulated cells remained viable for 1 week *in vitro* (Fig. 3E, 3F) and secreted PGT121 at up to 3.60 µg per device per day in devices loaded with 8 × 10^6^ cells (Fig. 3G). H&E-stained cross-sections of devices loaded with 8 × 10^6^ MSCs after 1 week of *in vitro* culture revealed early remodeling of the biodegradable matrix by the cells, with no evidence of cell escape through the nanofibrous membrane (Fig. 3H). Taken together, these results establish that clonally selected, high-producing MSCs retain viability and secretory function following encapsulation, and that PGT121 readily diffuses across the ZPU nanofibrous membrane.

### *In vivo* evaluation of encapsulated cell survival and therapeutic protein delivery across multiple mouse models

The first implantation study focused on cell survival. GFP/Luc-MSCs were encapsulated into ZPU devices and implanted into the dorsal subcutaneous space of BALB/c mice. BLI confirmed sustained cell survival over 16 weeks post-implantation (Fig. S4A, S4B). The device maintained structural integrity with no evidence of deformation (Fig. S4C). Upon retrieval at 16 weeks, devices displayed intact photocurable sealing, evident vascularization around the device surface (Fig. S4D), and positive bioluminescence signal (Fig. S4E). H&E-stained (Fig. S4F) and anti-GFP-stained (Fig. S4G) cross-sectional images of retrieved devices confirmed no evidence of encapsulated cell escape or host immune cell infiltration through the nanofibrous membrane, with cells uniformly distributed throughout the device lumen. These findings demonstrate that the encapsulation device supports long-term cell survival in the subcutaneous space of a fully allogeneic, immunocompetent host.

With cell survival confirmed, we next assessed *in vivo* therapeutic protein production and systemic delivery. As a human antibody, PGT121 elicits anti-drug antibody (ADA) responses that neutralize circulating PGT121 and accelerate its clearance in xenogeneic hosts. To avoid ADA interference with the assessment of PGT121 production from the device, we selected RAG2-knockout (RAG2-KO) mice, which lack functional T and B cells but retain intact innate immunity. In the second implantation study, devices loaded with PGT121/GFP/Luc-MSCs were implanted into the dorsal subcutaneous space of RAG2-KO mice. Serum PGT121 concentrations remained above 5 µg mL^−1^ for most of the 36-week implantation period until device retrieval (Fig. 4A). Devices retrieved at 8 (Fig. S5A), 16, and 36 weeks (Fig. 4B) maintained structural integrity and developed extensive per-device vasculature. BLI of retrieved devices showed uniform signals along the device length at 8, 16, and 36 weeks, indicating sustained cell viability throughout the implantation period (Fig. 4C, 4D). *Ex vivo* culture of devices demonstrated continued PGT121 secretion of approximately 3.5, 4.0, and 2.6 µg device^−1^ day^−1^ for devices retrieved at 8, 16, and 36 weeks, respectively (Fig. 4E), confirming maintenance of production capacity after long-term implantation. H&E-stained cross-sectional images of retrieved devices at 16 (Fig. S5B) and 36 weeks (Fig. 4F) showed a thin fibrotic layer surrounding the device, no host immune cell infiltration through the nanofibrous membrane, and vascular lumens containing erythrocytes in the peri-device tissue. Anti-GFP immunostaining showed GFP-positive cells confined to the device lumen with none detected in surrounding tissue at both 16 (Fig. S5C) and 36 weeks (Fig. 4G), confirming the absence of cell escape through the nanofibrous membrane.

**Fig. 4.**
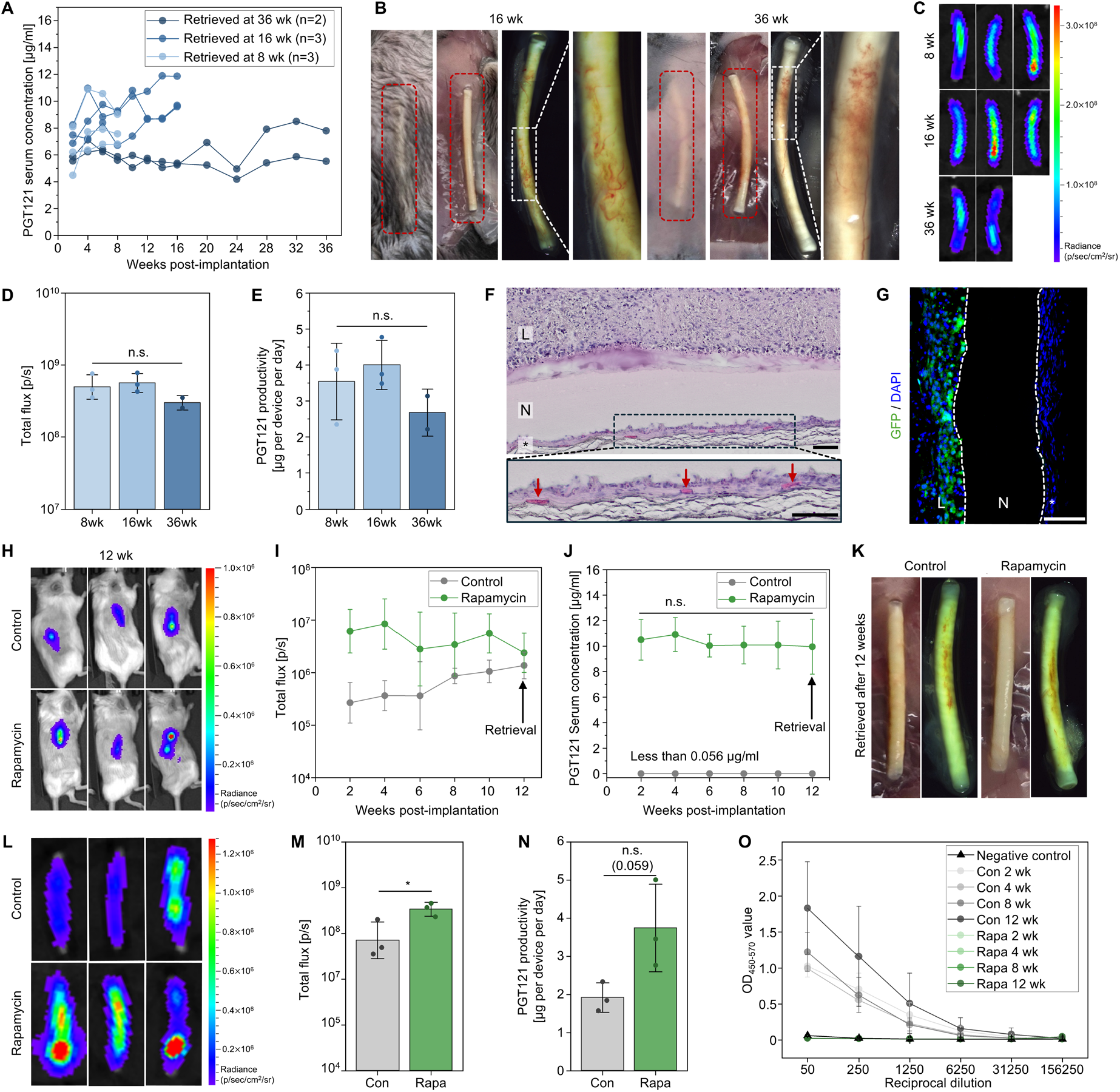
*In vivo* evaluation of encapsulated cell survival and therapeutic protein delivery across multiple mouse models. **(A-H)** Devices loaded with PGT121/GFP/Luc-MSCs were implanted into the dorsal subcutaneous space of RAG2-KO mice. (*n = 8* up to 8 weeks, *n = 5* up to 16 weeks, and *n = 2* up to 36 weeks) **(A)** Serum PGT121 concentrations measured biweekly up to 16 weeks and at 4-week intervals thereafter up to 36 weeks post-implantation **(B)** Representative images of the implantation site and retrieved devices at 16 weeks (left panel) and 36 weeks (right panel). Left to right within each panel: implantation site (dorsum), exposed devices after skin incision, optical micrograph of a retrieved device, and magnified view of peri-device vascularization. **(C)** Bioluminescence images of retrieved devices at 8, 16, and 36 weeks (top to bottom) and **(D)** corresponding quantification. **(E)** *Ex vivo* per-device PGT121 productivity of devices retrieved at 8, 16, and 36 weeks. n.s., not significant among all groups (p > 0.05). **(F)** H&E-stained cross-sectional images of a device retrieved at 36 weeks. Arrows indicate vascular lumens. Scale bar, 100 µm. **(G)** Immunofluorescence images of a device retrieved at 36 weeks. GFP (green) and DAPI (blue). White dashed lines indicate the boundary of the nanofibrous membrane. Scale bar, 100 µm. In **(F)** and **(G)**: L, device lumen; N, nanofibrous membrane; *, fibrotic layer. **(H–O)** Devices loaded with PGT121/GFP/Luc-MSCs were implanted into the dorsal subcutaneous space of BALB/c mice receiving either no treatment (*n = 3*) or systemic rapamycin (*n = 3*). **(H)** Bioluminescence images of control (top) and rapamycin-treated (bottom) mice at 12 weeks and **(I)** corresponding quantification up to device retrieval at 12 weeks. **(J)** Serum PGT121 concentrations measured biweekly up to 12 weeks. n.s., not significant across time points in the rapamycin-treated group (p > 0.05). **(K)** Representative images at 12 weeks. Control (left panel) and rapamycin-treated (right panel) groups. Within each panel, exposed devices after skin incision (left) and optical micrograph of a retrieved device (right) **(L)** Bioluminescence images of retrieved devices from the control (top) and rapamycin-treated (bottom) groups at 12 weeks and **(M)** corresponding quantification. *\*p* < 0.05. **(N)** *Ex vivo* per-device PGT121 productivity at 12 weeks. n.s., not significant. **(O)** Anti-PGT121 antibody levels measured from serially diluted serum; naive BALB/c mice served as negative control.

We next employed a complementary pharmacological approach to evaluate device performance in an immunocompetent allogeneic host. We used rapamycin, an mTOR inhibitor, to suppress ADA formation while preserving a partially attenuated but functional host immune response. In this third implantation study, devices loaded with PGT121/GFP/Luc-MSCs were implanted into the dorsal subcutaneous space of BALB/c mice with or without systemic rapamycin treatment. BLI confirmed sustained cell survival over 12 weeks, with higher average signal observed in the rapamycin-treated group, likely reflecting partially attenuated immune responses (Fig. 4H, 4I). Serum PGT121 concentrations in the rapamycin-treated group stabilized at ∼10 µg mL^−1^, whereas it remained undetectable in the untreated group (below the lower limit of detection, 0.056 µg mL^−1^) (Fig. 4J). Devices retrieved at 12 weeks from both groups remained structurally intact and were surrounded by well-developed vasculature (Fig. 4K). BLI (Fig. 4L, 4M) and *ex vivo* PGT121 secretion assays (Fig. 4N) of retrieved devices confirmed sustained cell survival and PGT121 production in both groups, again with higher levels observed in the rapamycin-treated group. The discrepancy between undetectable serum PGT121 and sustained *ex vivo* PGT121 production from retrieved devices in the untreated group corresponded with the detection of anti-PGT121 antibodies, observed only in this group (Fig. 4O), indicating that the absence of circulating PGT121 resulted from ADA-mediated clearance rather than loss of *in vivo* device function. Collectively, these results across three distinct mouse models demonstrate that the ZPU encapsulation device supports long-term survival and function of encapsulated cells in the subcutaneous space.

### *In vitro* and *in vivo* evaluation of cryopreserved cell-loaded devices for off-the-shelf use

We next evaluated whether cell-loaded devices could be cryopreserved for off-the-shelf use. SEM imaging confirmed that the nanofibrous ZPU membrane morphology was preserved after storage in liquid nitrogen followed by thawing (Fig. S6A). We then evaluated the viability and function of cell-loaded devices after the freeze-thaw cycle, both *in vitro* and *in vivo*, following the experimental timeline outlined in Fig. S6B. BLI of devices loaded with PGT121/GFP/Luc-MSCs before and after the freeze-thaw cycle confirmed uniform cell distribution throughout the devices, with ∼55% reduction in signal intensity after thawing (Fig 5A, 5B). Per-device PGT121 productivity was similarly reduced by ∼51% (Fig. 5C), consistent with the reduction in bioluminescence signal intensity, a commonly observed consequence of the cryopreservation process. H&E-stained cross-sections of thawed devices showed no cell escape or structural damage to the nanofibrous membrane after the freeze-thaw cycle (Fig. S6C). To validate *in vivo* performance, cryopreserved cell-loaded devices were thawed and implanted into the dorsal subcutaneous space of RAG2-KO mice. Serum PGT121 concentrations were maintained at ∼10 µg mL^−1^ over 12 weeks until device retrieval (Fig. 5D). Retrieved devices remained structurally intact and were surrounded by well-vascularized tissue (Fig. 5E). Bioluminescence signal intensity (Fig. 5F, 5G) and *ex vivo* PGT121 productivity (Fig. 5H) of retrieved devices were comparable to pre-implantation levels. H&E-stained (Fig. 5I) and anti-GFP-stained (Fig. 5J) cross-sectional images of retrieved devices confirmed that the immunoprotective function of the device were preserved after cryopreservation and long-term implantation, with cells uniformly distributed throughout the device lumen. H&E-stained sections (Fig. 5I) also showed vascular lumens containing erythrocytes in the peri-device tissue, indicating active vascularization around the device. Together, these results demonstrate that cell-loaded devices can be cryopreserved, and implanted without loss of therapeutic function, supporting their feasibility as off-the-shelf products for use in resource-limited settings.

**Fig. 5.**
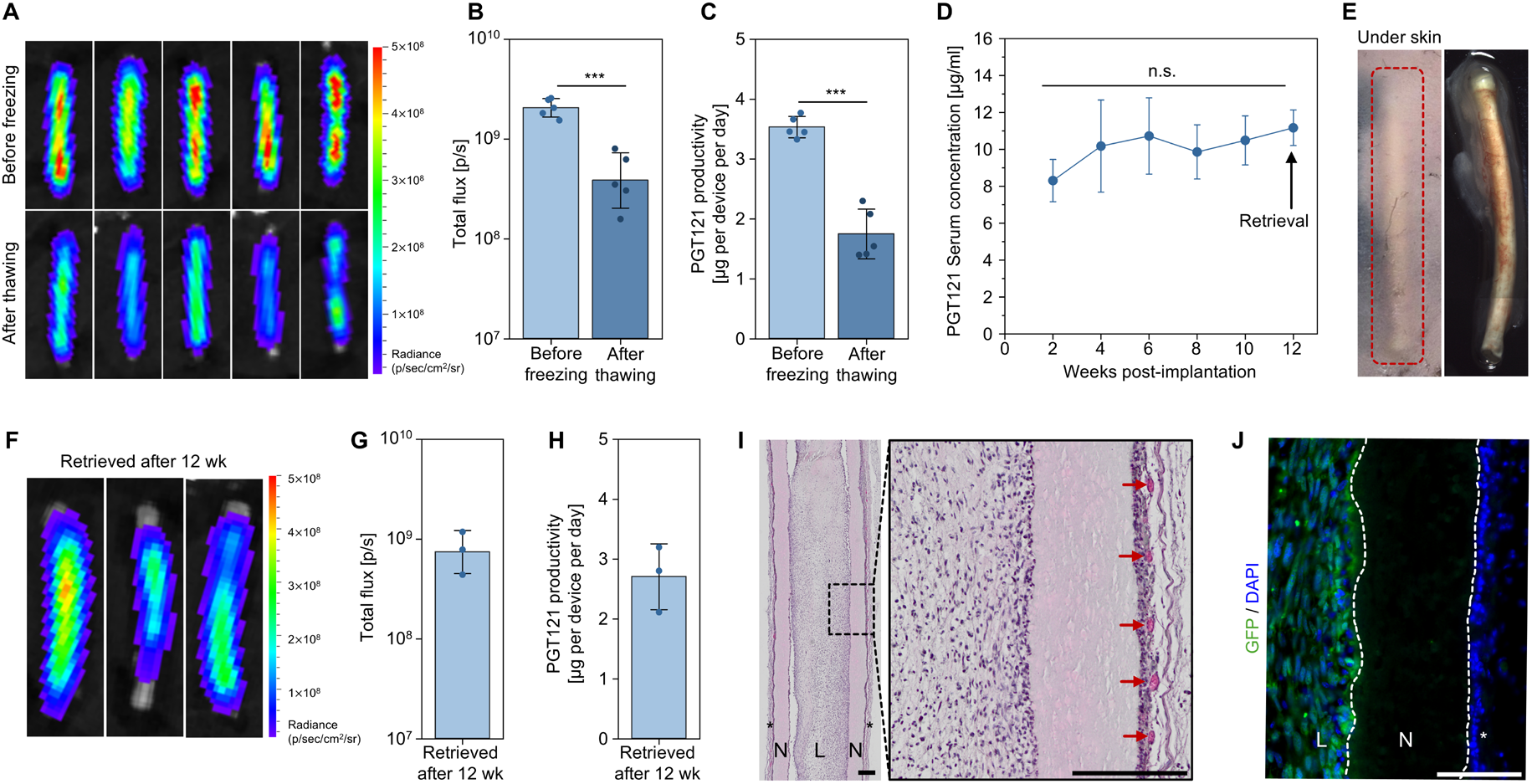
*In vitro* and *in vivo* evaluation of cryopreserved cell-loaded devices for off-the-shelf use. **(A)** Bioluminescence images of devices loaded with PGT121/GFP/Luc-MSCs before freezing (top) and after thawing (bottom), and **(B)** corresponding quantification. **(C)** *In vitro* per-device PGT121 productivity before freezing and after thawing. In **(B)** and **(C)**, *n = 5, ***p* < 0.001. **(D–J)** Cryopreserved cell-loaded devices were thawed and implanted into the dorsal subcutaneous space of RAG2-KO mice. **(D)** Serum PGT121 concentrations biweekly up to device retrieval at 12 weeks (*n = 3*). *n*.*s*., not significant across time points (p > 0.05). **(E)** Representative images at 12 weeks post-implantation. Left to right: implantation site (dorsum), exposed devices after skin incision, and optical micrograph of a retrieved device. **(F)** Bioluminescence images of retrieved devices at 12 weeks and **(G)** corresponding quantification. **(H)** *Ex vivo* per-device PGT121 productivity at 12 weeks. **(I)** H&E-stained cross-sectional images of a device retrieved at 12 weeks. Arrows indicate vascular lumens. Scale bar, 200 µm. **(J)** Immunofluorescence images of retrieved device at 12 weeks. GFP (green) and DAPI (blue). White dashed lines indicate the boundary of the nanofibrous membrane. Scale bar, 200 µm. In **(I)** and **(J)**: L, device lumen; N, nanofibrous membrane; *, fibrotic layer.

### *In vivo* evaluation of hiMSCs as a clinically relevant therapeutic cell source

Having established device performance using murine MSCs, we next evaluated a clinically relevant human cell source. We selected hiMSCs for their reduced donor heterogeneity, scalability, and enhanced proliferative capacity while retaining key MSC functional properties(*22*). First, to assess encapsulated cell survival in a xenogeneic setting, devices loaded with GFP/Luc-expressing hiMSCs were implanted into the dorsal subcutaneous space of immunocompetent BALB/c mice. BLI confirmed sustained cell survival over 12 weeks (Fig. 6A, 6B). Retrieved devices remained structurally intact (Fig. 6C) and retained positive bioluminescence signal (Fig. 6D). We then engineered hiMSCs to produce PGT121 via lentiviral transduction using the same workflow established for murine MSCs, achieving a PGT121 production rate of 0.87 µg per 10^6^ cells per day (Fig. 6E). HiMSC spheroids (Fig. S7A) were loaded into devices and cultured *in vitro*, and both cell survival (Fig. S7B, S7C) and PGT121 production (Fig. S7D) were confirmed following loading. For *in vivo* evaluation, devices loaded with PGT121/GFP/Luc-hiMSCs were implanted into the dorsal subcutaneous space of RAG2-KO mice. Serum PGT121 concentrations were maintained between 2 and 3 µg mL^−1^ over 12 weeks until device retrieval (Fig. 6F). Retrieved devices remained structurally intact (Fig. 6G, Fig. S7E, S7F). BLI (Fig. 6H, 6I) and *ex vivo* PGT121 productivity of approximately 0.27 µg device^−1^ day^−1^ (Fig. 6J) confirmed that encapsulated cells retained viability and PGT121 production capacity after long-term implantation. H&E-stained (Fig. 6K) and anti-GFP-stained (Fig. 6L) cross-sectional images of retrieved devices confirmed that the immunoprotective function of the device were preserved, with cells uniformly distributed throughout the device lumen. These results provide proof of concept extending the use of the ZPU encapsulation platform to hiMSCs, a clinically relevant human cell source, as a therapeutic cell type.

**Fig. 6.**
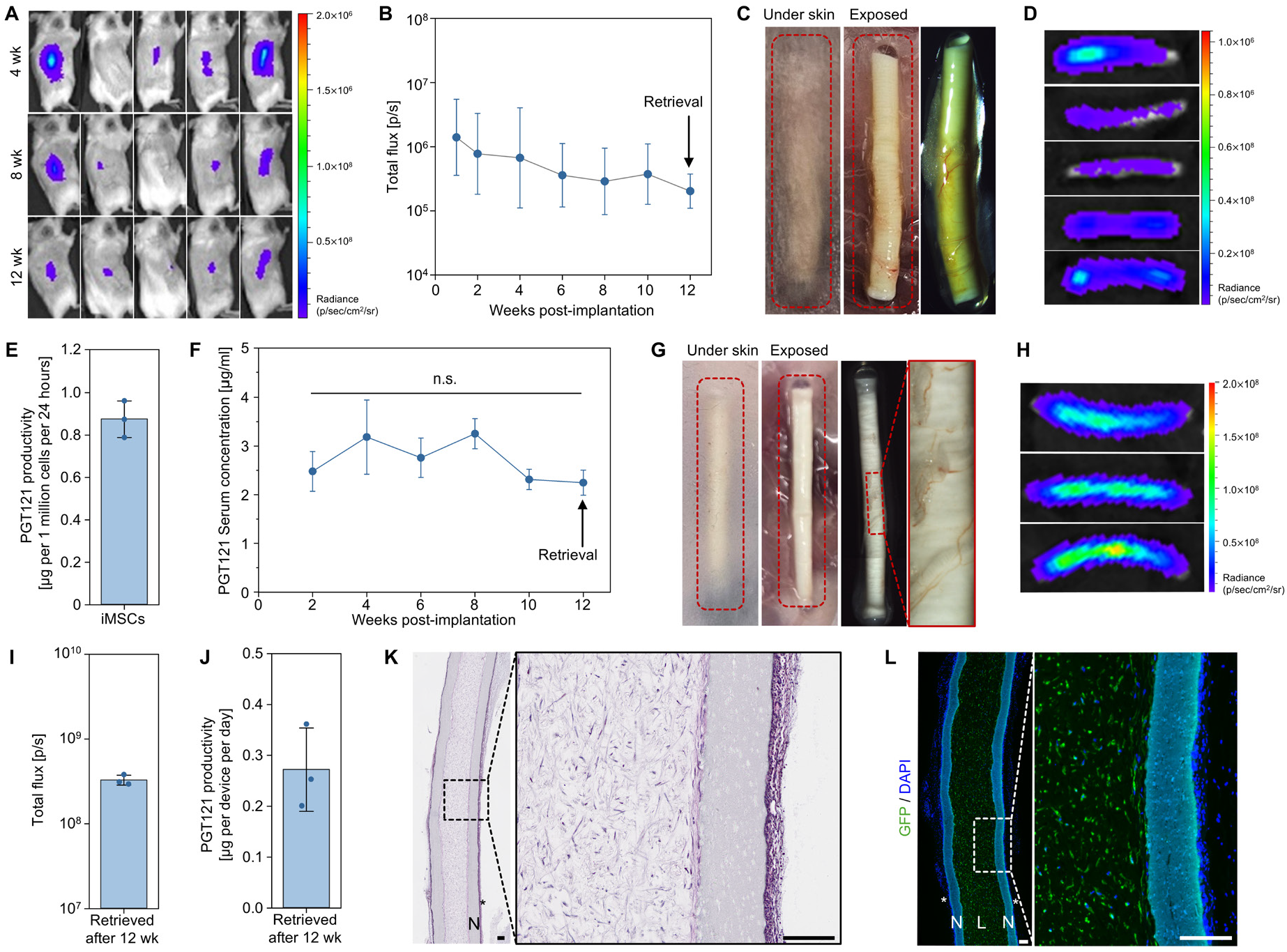
*In vivo* evaluation of hiMSCs as a clinically relevant therapeutic cell source. **(A–D)** Devices loaded with GFP/Luc-hiMSCs were implanted into the dorsal subcutaneous space of BALB/c mice (*n = 5*). **(A)** Bioluminescence images of mice with implanted devices at 4, 8, and 12 weeks post-implantation and **(B)** corresponding quantification measured biweekly up to device retrieval at 12 weeks. **(C)** Representative images at 12 weeks post-implantation. Left to right: photograph of the implantation site (dorsum), exposed devices after skin incision, and optical micrograph of a retrieved device. **(D)** Bioluminescence images of retrieved devices at 12 weeks post-implantation. **(E)** Per-cell PGT121 productivity of PGT121-producing hiMSCs (*n = 3*). **(F–L)** Devices loaded with PGT121/GFP/Luc-hiMSCs were implanted into the dorsal subcutaneous space of RAG2-KO mice. **(F)** Serum PGT121 concentrations measured biweekly up to device retrieval at 12 weeks (*n = 3*). n.s., not significant across time points (p > 0.05). **(G)** Representative images at 12 weeks post-implantation. Left to right: photograph of the implantation site (dorsum), exposed devices after skin incision, optical micrograph of a retrieved device, and magnified image showing peri-device vascularization. **(H)** Bioluminescence images of retrieved devices at 12 weeks and **(I)** corresponding quantification. **(J)** *Ex vivo* per-device PGT121 productivity at 12 weeks. **(K)** H&E-stained cross-sectional images of a device retrieved at 12 weeks. Scale bar, 200 µm. **(L)** Immunofluorescence images of a device retrieved at 12 weeks. GFP (green) and DAPI (blue). White dashed lines indicate the boundary of the nanofibrous membrane. Scale bar, 200 µm. In **(K)** and **(L)**: L, device lumen; N, nanofibrous membrane; *, fibrotic layer.

### Proof-of-concept evaluation of procedural feasibility and device function in a minipig model

We next conducted a proof-of-concept study in Göttingen minipigs to determine whether the procedural simplicity and device performance established in mice could be maintained at a clinically relevant scale. Porcine skin and subcutaneous tissue closely resemble those of humans in epidermal thickness, dermal collagen composition, and vascular anatomy, as well as in immune system characteristics (*23*), enabling evaluation at a clinically relevant scale. Devices loaded with PGT121/GFP/Luc-expressing murine MSCs were implanted into the abdominal subcutaneous space of Göttingen minipigs. A scaled-up applicator with an enlarged body and a longer needle was used to facilitate handling in large animals and accommodate longer devices. For the minipig study, devices with the same inner diameter as in the mouse study (1.2 mm) but with an extended length (4 cm), comparable in scale to a contraceptive implant, were used. The implantation procedure followed three sequential steps: positioning of the device-loaded needle with the slider locked, insertion into the subcutaneous space, and device deployment by unlocking the slider and retracting the needle while the obturator secured the device at the target site (Fig. 7A). For retrieval, some devices were retrieved together with the surrounding tissue for histological analysis, while others were retrieved using a clinically relevant procedure: a skin incision was made, the fibrotic layer was partially dissected at one end to expose the device tip, and the device was gently slid out (Fig. S8C). Devices retrieved by slide-out showed a clean surface free of tissue adhesion (Fig. 7B, left), whereas devices retrieved with the surrounding tissue retained the attached fibrotic layer as expected (Fig. 7B, middle and right). In both cases, devices maintained structural integrity without deformation or damage. Despite the vigorous xenogeneic immune response expected from mouse-to-pig implantation, encapsulated cells retained detectable bioluminescence signals (Fig. 7C, 7D) and sustained PGT121 production (Fig. 7E), although at substantially reduced levels compared to the murine studies. H&E-stained cross-sections of devices from both retrieval approaches revealed no cell escape or host immune cell infiltration through the nanofibrous membrane, minimal residual fibrotic layer on slide-out–retrieved devices (Fig. 7F). Taken together, these results demonstrate the feasibility of the encapsulation platform in a large-animal model, with minimally invasive implantation and retrieval procedures, and preserved structural integrity and immunoprotective function in the porcine subcutaneous space.

**Fig. 7.**
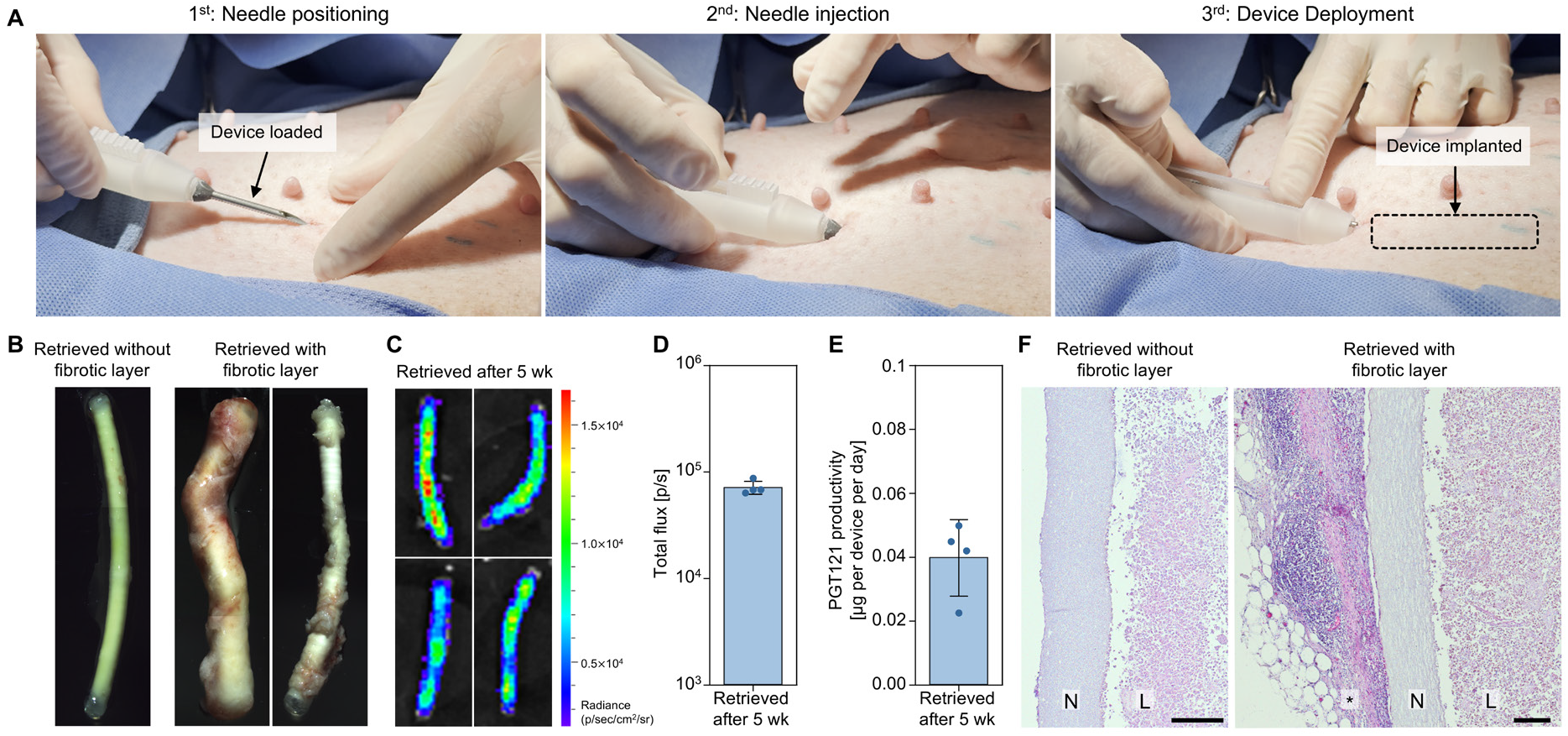
Proof-of-concept evaluation of procedural feasibility and device function in a minipig model. **(A)** Subcutaneous implantation procedure using the scaled-up applicator. Sequential steps (left to right): needle positioning, needle insertion, and device deployment. **(B–F)** Devices loaded with PGT121/GFP/Luc-murine MSCs were implanted into the abdominal subcutaneous space of Göttingen minipigs (*n = 4*). **(B)** Representative optical micrographs of retrieved devices at 5 weeks post-implantation. Left: device retrieved by slide-out after partial dissection of the fibrotic layer. Middle and right: device retrieved with the surrounding fibrotic layer intact. **(C)** Bioluminescence images of retrieved devices at 5 weeks and **(D)** corresponding quantification. **(E)** *Ex vivo* per-device PGT121 productivity at 5 weeks. **(F)** H&E-stained cross-sectional images of devices retrieved by slide-out (left) and retrieved with the fibrotic layer intact (right) at 5 weeks. L, device lumen; N, nanofibrous membrane; *, fibrotic layer. Scale bar, 200 µm.

## DISCUSSION

Cell encapsulation has been pursued as a strategy to achieve sustained therapeutic protein delivery through immunoprotected implantable cellular factories, potentially eliminating the need for repeated injections. Multiple encapsulation devices have been evaluated in clinical trials, predominantly for islet implantation in type 1 diabetes (PEC-Encap, NCT02239354; βAir, NCT02064309; VX-264, NCT05791201)(*10, 24*), yet long-term therapeutic efficacy has been limited, with fibrotic reactions and inadequate mass transfer consistently identified as the primary barriers to sustained device performance. The recent FDA approval of Encelto, the first approved genetically engineered encapsulated cell therapy, demonstrates that the platform can succeed clinically(*25*). However, its delivery requirements represent a considerably less demanding setting than systemic protein delivery: the implant delivers nanograms per day of protein directly to its target cells within the immune-privileged vitreous cavity, where local retention sustains therapeutic concentrations(*8*). By contrast, most other therapeutic applications of cell encapsulation require implantation at clinically relevant but non-immune-privileged sites that elicit pronounced FBR, and demand substantially higher sustained systemic concentrations. For example, islet transplantation for type 1 diabetes requires a cumulative dose on the order of 10^9^ cells to achieve sustained insulin independence, and delivery of broadly neutralizing antibodies against HIV-1, such as PGT121 as employed in this study, necessitates sustained serum concentrations of approximately 10 µg mL^−1^ for therapeutic efficacy(*26, 27*). Collectively, these requirements underscore the need for technical advances that overcome fibrotic reaction and mass transfer barriers to enable broader clinical translation of cell encapsulation.

In this study, we address these challenges through the integration of optimized device design and cell engineering strategies. First, surface chemistry is a critical determinant of the FBR and thus directly impacts long-term device performance(*28*). Zwitterionic polymers, which resist nonspecific protein adsorption, have emerged as a promising class of antifouling biomaterials and are being actively investigated in both preclinical and clinical settings for implantable medical devices(*29*). We fabricated the encapsulation membrane from a custom-synthesized ZPU to minimize material-driven FBR and maintain facile mass transfer. We employed electrospinning for the encapsulation membrane and systematically optimized key electrospinning parameters to achieve the mechanical robustness required for long-term subcutaneous implantation. Another key design consideration for subcutaneous cell encapsulation devices is the form factor. Subcutaneous contraceptive implants such as Nexplanon, although non-cellular hormone-delivery devices, provide an established clinical precedent as a slender cylindrical rod that is inserted via a simple applicator-based procedure (*14*). Inspired by the widespread clinical adoption of this form factor, including particularly rapid uptake in some less developed regions of the world, we adopted a cylindrical geometry enabling minimal invasives procedures without specialized surgical facilities. Finally, the choice of matrix co-encapsulated with the cells is an important consideration. Alginate has been widely used in islet encapsulation for its biocompatibility and non-degradable immunoprotecting properties(*30*). However, in our device, the nanofibrous membrane already provides immunoprotection, allowing the matrix to serve primarily as a support for cell survival and function. We employed a biodegradable fibrin-collagen blended matrix that supports initial cell attachment and is gradually degraded as MSCs proliferate to reach a device capacity governed by mass transfer equilibrium. Consistent with this design rationale, devices retrieved after up to 36 weeks of subcutaneous implantation maintained structural integrity with thin fibrotic layers, no cell escape or immune cell infiltration, uniform cell distribution throughout device lumen, and active peri-device vascularization.

Beyond device design, the choice of cell type as a therapeutic protein producer is a critical consideration. Key requirements include amenability to stable genetic modification, robust *in vitro* expansion capacity, and an established safety profile. MSCs not only meet these criteria but also offer immunomodulatory capacity, low immunogenicity, and pro-angiogenic properties, with over 1,000 clinical trials registered across diverse therapeutic areas and recent FDA approval of the first allogeneic MSC therapy(*19, 31*). For our murine studies, we used murine MSCs to model an allogeneic implantation setting. MSCs were engineered to constitutively produce PGT121 using a third-generation self-inactivating lentiviral vector, a platform that has enabled multiple FDA-approved CAR-T cell and hematopoietic stem cell gene therapies(*32*). For our proof-of-concept studies with human cells, we selected hiMSCs, which are increasingly being evaluated in clinical trials. Although primary human MSCs would be the most direct counterpart to the murine MSCs used in this study, hiMSCs retain the core advantages of human MSCs while offering a rejuvenated proliferative phenotype, reduced heterogeneity through single-clone iPSC derivation, and the capacity for scalable expansion from a single master cell bank(*31*).

The choice of *in vivo* model is another key consideration for evaluating the combined performance of an optimized device and engineered cells. Subcutaneous implantation of allogeneic cell encapsulation devices poses three principal challenges: (i) limited oxygen and nutrient availability inherent to the subcutaneous space, (ii) FBR-induced fibrotic layer formation that further restricts mass transfer, and (iii) allogeneic immune rejection. We first selected a fully allogeneic model, transplanting C57BL/6-derived MSCs into BALB/c mice, to evaluate cell survival in the presence of all three challenges. Then, to evaluate therapeutic protein production and systemic delivery, we employed two complementary mouse models, each offering a distinct strategy to address a key complicating factor in our preclinical setting: the immunogenicity of human PGT121 in xenogeneic hosts, which elicits ADA responses that complicate the assessment of therapeutic protein production(*33*). The RAG2-KO model, which lacks functional adaptive immunity but retains intact innate immunity, enabled assessment of therapeutic protein delivery in the absence of ADA interference while largely preserving the FBR, as fibrotic layer formation is primarily driven by macrophages; Doloff et al. reported that fibrotic reactions against biomaterial implants in RAG2-KO mice were comparable to that in wild-type C57BL/6 mice(*34*). Thus, RAG2-KO mice present challenges (i) and (ii) albeit in the absence of challenge (iii). The rapamycin-treated immunocompetent BALB/c model provided a complementary pharmacological approach to suppress ADA formation. Rapamycin, an mTOR inhibitor with established use in solid organ transplantation, has also been shown to mitigate ADA formation by suppressing B cell differentiation and promoting regulatory T cell expansion(*35, 36*). While rapamycin partially attenuates allogeneic rejection and may also reduce the fibrotic reactions, fibrotic layer formation is primarily macrophage-driven and not directly targeted by mTOR inhibition(*34, 37*), and is therefore unlikely to be fully eliminated in this model. Thus, this rapamycin-treated allogeneic model complements the RAG2-KO study by providing an alternative approach to avoiding ADA interference. The sustained therapeutic protein delivery observed across both models indicates that the device supports long-term cell survival and protein production. Importantly, the ADA response observed in this study is specific to the murine setting, where a fully human antibody is produced in a non-human host, and is unlikely to be a significant barrier in human applications, as minimal ADA induction has been reported in clinical trials of PGT121(*15*).

From a translational perspective, two findings support the feasibility of this platform in clinical settings with limited infrastructure. First, the manufacturing infrastructure for gene-modified cell products remains concentrated in high-income countries, even though diseases such as HIV, where over two-thirds of affected individuals reside in sub-Saharan Africa, disproportionately burden low- and middle-income countries (*38, 39*). Our platform addresses this gap through a cryopreservation-based off-the-shelf approach that decouples device manufacturing from the point of care. Cell-loaded devices retained therapeutic function after cryopreservation and thawing, validating a workflow of centralized manufacturing and quality control, followed by on-site implantation. Second, the procedural simplicity of our platform further supports its use in settings without surgical facilities. Existing cell encapsulation devices evaluated in clinical trials require surgical implantation and retrieval, and complications related to these procedures have been among the most frequently reported adverse events (27.9% in NCT03163511)(*24*). In this study, devices were successfully implanted into the abdominal subcutaneous space of Göttingen minipigs using a scaled-up applicator without sutures for wound closure. Furthermore, the antifouling properties of the ZPU surface enabled retrieval through a slide-out procedure without tissue adhesion. These procedures are comparable to those of subcutaneous contraceptive implants, which are routinely performed by trained healthcare workers across sub-Saharan Africa(*14*). Together, these approaches lower the barrier to adoption in resource-limited settings, providing a pathway toward clinical translation.

Despite the promising features of our therapeutic cell encapsulation platform, several limitations should be acknowledged. First, per-cell PGT121 productivity decreased substantially after encapsulation, likely reflecting metabolic constraints imposed by high cell density within the confined device lumen, consistent with previous reports (*40, 41*). Although serum PGT121 concentrations were sustained at µg mL^−1^ levels in our murine models, clinical translation to humans may demand output that exceeds the current capacity of a single device. Future studies aimed at improving per-device productivity through integrative strategies including cell-supporting scaffolds and screening of alternative cell types such as ARPE-19(*42*) could help achieve therapeutic concentrations in larger hosts. Second, although sustained serum PGT121 concentrations were demonstrated across multiple murine models, this study did not evaluate therapeutic efficacy in a disease model. Future studies should assess therapeutic outcomes in relevant disease models to demonstrate a direct link between sustained protein delivery and therapeutic efficacy. Third, cell viability and protein production in the minipig model were substantially reduced compared to the murine studies, most likely reflecting the vigorous xenogeneic immune response arising from mouse-to-pig implantation. This barrier would be absent in a species-matched setting, and future large-animal studies using species-matched cells would more accurately reflect the anticipated clinical scenario. As a future direction, engineering cells to co-express multiple therapeutic proteins, or co-loading distinct cell populations, could provide a versatile approach to enhance device performance or enable addressing multiple therapeutic targets from a single implant.

In summary, we developed and evaluated a miniaturized subcutaneous cell encapsulation platform to enable sustained therapeutic protein delivery, with particular emphasis on accessibility in resource-limited settings. We demonstrated sustained cell survival and therapeutic protein production in mice, off-the-shelf potential through cryopreservation, and applicator-based implantation and retrieval at clinically relevant scale in minipigs. Collectively, these results establish a versatile encapsulation platform as a translatable strategy for sustained biologic delivery in resource-limited clinical settings.

## MATERIALS AND METHODS

### Study design

The objective of this study was to design a miniaturized subcutaneous cellular implant for sustained therapeutic protein delivery in resource-limited settings. We assessed cell viability and sustained therapeutic protein production following subcutaneous implantation of cell-loaded devices in multiple murine models. Murine MSCs were evaluated in three complementary mouse models: (i) immunocompetent allogeneic mice for cell survival assessment, (ii) immunocompromised RAG2-KO mice for PGT121 production assessment, and (iii) rapamycin-treated immunocompetent mice for PGT121 production assessment under partially attenuated host immunity. Human iMSCs were similarly evaluated in immunocompetent xenogeneic hosts for cell survival assessment and in immunocompromised mice for PGT121 production assessment. Procedural compatibility of the platform at clinically relevant scale was evaluated in Göttingen minipigs as a proof-of-concept large-animal study. The number of cells loaded per device was determined on the basis of *in vitro* production data and prior encapsulation experience. Sample sizes were chosen on the basis of historical data to ensure adequate statistical power and are specified in the corresponding figure legends. All mice used were females to eliminate any potential confounding influences of gender differences. Mice were randomly assigned to treatment groups. No animals or data points were excluded from analysis. The number of biological replicates is specified in each figure legend. All animal procedures were performed in accordance with protocols approved by the Cornell Institutional Animal Care and Use Committee (IACUC).

### Fabrication of zwitterionic polyurethane nanofibrous membranes via electrospinning

ZPU polymers were dissolved in HFIP at concentrations of 10, 12.5, 15, 17.5, and 20% (w/v) under stirring at room temperature. The solution was loaded into a 20 mL plastic syringe (Norm-Ject) and dispensed at 1.2 mL h^−1^ via a syringe pump (Harvard Apparatus). Electrospinning was performed at 18 kV through a 23-gauge blunt needle with a tip-to-collector distance of 12 cm. Aluminum rods of varying dimensions (McMaster-Carr) were coated with sucrose solution (∼1.07 g mL^−1^) and used as rotating collectors (400–450 rpm). After 45 min (Ver.1) and 90 min (Ver.2) of electrospinning, nanofibrous membranes were detached from the rods in a DI water bath, soaked in DI water overnight, and dried at room temperature. For devices to be sealed with a photocurable adhesive (1187-M-SV01, Dymax) required pretreatment prior to cell loading, as direct application to wet surfaces after cell loading resulted in poor adhesion and seal detachment. Dried membranes were exposed to air plasma for 60 s (Harrick Plasma), then coated with the adhesive on both inner and outer end surfaces and cured under a UV lamp (365 nm, 60 W). The tubes were sterilized with ethylene oxide (Anprolene AN74, Anderson Sterilizers) and stored until use.

### Encapsulation of cells into zwitterionic polyurethane nanofibrous device

Spheroids prepared the day before were collected into 1mL microcentrifuge tube and resuspended in 30 µL of the desired matrix. For alginate-based devices, 30 μL of 2% (w/v) SLG100 alginate solution dissolved in 0.9% (w/v) NaCl was mixed with the spheroids and loaded into the device. The device was then submerged in a cross-linking buffer containing 100 mM CaCl_2_ and 5 mM BaCl_2_ for 10 min, followed by washing with 0.9% (w/v) NaCl solution. For degradable hydrogel-based devices, 28.5 μL of the prepared matrix (see Supplementary Materials) was mixed with spheroids, and 1.5 μL of thrombin (100 U mL^−1^ stock; 5 U mL^−1^ final) was added immediately before loading into the device. The resulting 30 μL of cell–matrix mixture was then loaded into the device. For thermal sealing, the open ends of the membrane were sealed using a hand impulse sealer. For photocurable adhesive sealing, membranes pretreated with photocurable adhesive (as described above) were used. The adhesive was applied as a droplet to cap the open ends, then cured by exposure to a UV lamp (365 nm, 60 W) for 60 s. Sealed devices were maintained in culture medium at 37 °C until use. To confirm cell distribution within the devices, a small section was cut from each end, and the outer membrane layer was peeled off. The exposed surface was then examined under an optical microscope (EVOS AMF4300). For GFP-expressing cells, fluorescence imaging was additionally performed.

### Device transplantation with a custom-designed applicator and device retrieval in mice

A custom-designed applicator was fabricated by 3D printing (Form 2, Formlabs) and assembled with a 14-gauge needle (Fig. 2J). For transplantation, mice were anesthetized with 3% isoflurane in O_2_, and the dorsal skin was shaved and sterilized with betadine followed by 70% ethanol. The device was loaded into the needle of the assembled applicator, and the slider was locked into the notch on the applicator body. The needle was then inserted into the dorsal subcutaneous space. The slider was unlocked and the applicator needle was retracted to deploy the device at the target site, while the obturator secured the device in position, as described in Fig. 2K. The skin incision was closed with a wound clip. For retrieval, devices were collected following euthanasia by CO_2_ inhalation for histological analysis. Photographs were taken before and after skin incision to assess device integrity. Retrieval was also performed in live mice under anesthesia to demonstrate minimally invasive retrievability: a minor skin incision was made, the fibrotic layer was partially dissected to expose the device, and the device was gently slid out, as described in Fig. 2L. Device morphology upon retrieval was examined under a stereomicroscope (Olympus SZ61).

### Cell engineering

For generation of GFP/Luc-expressing cells, a pLenti-based expression vector encoding GFP and humanized firefly luciferase (Luc2) was constructed by VectorBuilder. For generation of PGT121-producing cells, a pLenti-based expression vector encoding a codon-optimized PGT121 with a signal peptide was constructed by VectorBuilder. The construct encodes PGT121 VH–CH (human IgG1 isotype) and PGT121 Vλ–Cλ linked by a T2A self-cleaving peptide, under the control of the EF1α promoter (Fig. S3A). Upon translation, the T2A linker enables cleavage of the heavy and light chains from a single transcript, which subsequently assemble into full-length PGT121. Lentiviral particles were produced by transfecting the transfer vector and packaging plasmids into 80–90% confluent HEK293T cells using the ViraPower Bsd Lentiviral Support Kit (Thermo Fisher Scientific, K497000). At 48 h post-transfection, the supernatant was collected, centrifuged to remove cell debris, and stored at −80 °C. For transduction, target cells were seeded in a six-well plates (5,000 cells per well), cultured overnight, and treated with 2 mL of virus-containing supernatant. After 24 h, the medium was replaced with fresh culture medium. For antibiotic selection of stably transduced cells, transduced cells were cultured in medium supplemented with 5 µg mL^−1^ blasticidin (PGT121) or 2 µg mL^−1^ puromycin (GFP/Luc) for 10 days. To establish high-producing clonal cell lines, single cells were sorted into 96-well plates using a FACSMelody (BD Biosciences). CloneR−2 (100-0691, STEMCELL Technologies) was supplemented according to the manufacturer’s instructions to promote clonal outgrowth. After 2 weeks of expansion, the 10 highest PGT121-producing clones were identified by primary screening (Fig. S3D), individually expanded, and reassessed in a secondary screening (Fig. 3B).

### Bioluminescence imaging (BLI)

To monitor engraftment and survival of encapsulated cells *in vivo*, mice with transplanted devices loaded with Luc-expressing cells were injected intraperitoneally (i.p.) with D-luciferin (150 mg kg^−1^ body weight; 14681, Cayman Chemical). For BLI of devices in *in vitro* or *ex vivo* experiments, devices were placed in a six-well plate containing 5 mL of D-luciferin solution (150 µg mL^−1^) in culture medium. BLI was performed using an IVIS Spectrum system (PerkinElmer) at the Biotechnology Resource Center at Cornell University. Regions of interest (ROIs) of identical size were applied to each sample, and total flux (photons s^−1^) was quantified using Living Image software (PerkinElmer).

### PGT121 quantification

To measure per-cell PGT121 productivity *in vitro*, 5 × 10^5^ cells were seeded in a T75 flask in 10 mL of culture medium, and conditioned medium was collected 24 h later. To measure per-device productivity, devices from *in vitro* studies or those retrieved after *in vivo* transplantation were incubated in 8 mL of culture medium for 24 h. All conditioned medium from cell culture and device incubation was centrifuged at 300 × *g* for 5 min to remove cell debris, and the supernatant was stored at −80 °C until analysis. To monitor serum PGT121 concentrations following transplantation, blood (approximately 100 μL) was collected from mice via retro-orbital bleeding at designated time points. Blood was collected into microcentrifuge tubes, allowed to clot at room temperature, and centrifuged at 1,800 × *g* for 15 min at 4 °C. Serum was separated, aliquoted, and stored at −80 °C until analysis.

PGT121 concentrations were quantified by antigen-capture ELISA. High-binding 96-well microplates (3590, Corning) were coated overnight at 4 °C with 100 μL per well of HIV-1 JRFL gp140 recombinant protein (B.JRFL gp140CF; ARP-12573, BEI Resources) at 1 μg mL^−1^ in coating buffer (421701-BL, BioLegend). Wells were blocked with 3% bovine serum albumin (BSA) in PBS for 1 h at room temperature (RT) and washed four times with wash buffer (PBS containing 0.05% Tween-20). Serially diluted PGT121 reference standard (ARP-12343, BEI Resources) and appropriately diluted samples were added at 100 μL per well and incubated for 2 h at RT with shaking. After four washes, horseradish peroxidase (HRP)-conjugated goat anti-human IgG Fc antibody (109-035-170, Jackson ImmunoResearch), diluted 1:5,000 in blocking buffer, was added and incubated for 1 h at RT with shaking. Wells were washed four times, and 100 μL of TMB substrate (34028, Thermo Fisher Scientific) was added. The reaction was stopped after 10 min by the addition of 100 μL of 2 N H_2_SO_4_, and absorbance was measured at 450 nm with background subtraction at 570 nm using a microplate reader. PGT121 productivity was calculated based on the measured concentration, with appropriate normalization to cell number or culture volume.

### Pig experiment

A scaled-up applicator was fabricated by 3D printing (Form 2, Formlabs). Göttingen minipigs were anesthetized with isoflurane and oxygen and maintained under general anesthesia throughout the procedure. The abdominal area was shaved and prepared under sterile conditions. Devices were loaded into the applicator and deployed into the abdominal subcutaneous space as described in Fig. 7A. The needle insertion site was covered with a Tegaderm transparent dressing (3M). For retrieval, a skin incision was made over the implantation site. Some devices were retrieved together with the surrounding tissue for histological analysis, while others were retrieved by slide-out after partial dissection of the fibrotic layer as described in Fig. S8C. Analysis of retrieved devices, including *ex vivo* PGT121 productivity, bioluminescence imaging, and histology, was performed as described above.

### Statistical analysis

Data are expressed as mean ± s.d. For quantification of bioluminescence signal, data were log-transformed prior to analysis and are presented as geometric mean ± geometric s.d. on a logarithmic scale. Comparisons between two groups were performed using unpaired, two-tailed Student’s *t*-tests. Comparisons among three or more groups were performed by one-way analysis of variance (ANOVA) followed by Tukey’s post hoc test. Sample sizes, including the number of mice per group, were chosen to ensure adequate statistical power based on historical data. No data were excluded from the analyses. All statistical analyses were performed using Origin software (OriginLab). Significance levels are denoted as n.s. (*P* ≥ 0.05), * (*P* < 0.05), ** (*P* < 0.01), and *** (*P* < 0.001). The number of biological replicates is specified in the figure legends.

## List of Supplementary Materials

Materials and Methods

Fig. S1 to S8

## Acknowledgements

We thank Cornell Center for Materials Research Facility for SEM imaging. We thank the Imaging Facility (RRID:SCR_021741) of the Biotechnology Resource Center of Cornell Institute of Biotechnology for their help with Bioluminescence imaging. All schematic figures were created using BioRender.

## Author contribution

M.L. and M.M. conceived and designed the project. B.W. designed lentiviral vector construct for engineering cells. K.W. and M.L. synthesized and characterized the ZPU polymer. A.B. designed the biodegradable matrix. K.O and M.L. designed the custom-designed applicator. M.L. performed all the other *in vitro* and mice experiments. J.A.F., K.O., and M.L. conducted the pig experiments. M.L. and M.M. wrote the manuscript. J.M.M.-M. discussed the data and revised the manuscript. All authors edited and reviewed the manuscript.

## Data and materials availability

All data associated with this study are presented in the paper and/or the Supplementary Materials.

