## Supplementary Materials for "Miniaturized subcutaneous cellular implants for sustained therapeutic protein delivery in resource-limited settings"

**List of Supplementary Materials**

Materials and Methods

Fig. S1 to S8

**Supplementary Materials:**

### Materials and methods

#### Materials

Sterile sodium alginate (Pronova, SLG100) was purchased from NovaMatrix. Calcium chloride dihydrate (C3306), barium chloride dihydrate (217565), poly(tetramethylene ether)glycol (PTMG, 345326), 1,4-diaminobutane (D13208), 1,1,1,3,3,3-hexafluoro-2-propanol (HFIP), fibrinogen (F8630), fibronectin (F2006), and thrombin (T4648) were purchased from Sigma-Aldrich. Collagen type I (354249) was purchased from Corning. Ultrapure water (10977015), Medium 199, Earle's Salts (10×) (21180021),1 M HEPES (1M) (15630080), and KnockOut™ serum replacement (10828028) were purchased from Thermo Fisher Scientific. N-butyldiethanolamine (B0725), 1,3-propanesultone (P0324), and 1,6-diisocyanatohexane (HDI, H0324) were purchased from TCI. Dichloromethane (D143SK-4) and sucrose (S5) were purchased from Fisher Scientific. Dimethyl sulfoxide (DMSO) was purchased from J.T. Baker. Stannous octoate ($Sn(Oct)_2$, 018590.22) was purchased from Alfa Aesar.

#### Animals

Eight-week-old female C57BL/6J, BALB/cJ, and B6.Cg-$Rag2^{tm1.1Cgn}$/J (RAG2-KO) mice were purchased from The Jackson Laboratory (Bar Harbor, ME). Göttingen minipigs (18 months old, 30 kg) were obtained from Marshall BioResources, North Rose NY. All animal procedures were approved by the Cornell Institutional Animal Care and Use Committee and complied with all relevant ethical regulations.

#### Cell culture

C57BL/6 mouse bone marrow mesenchymal stromal cells (MUBMX-01001) were purchased from Cyagen. Human iPSC-derived mesenchymal stromal cells (hiMSCs, #200-0781) were purchased from STEMCELL Technologies. HEK293T cell line (CRL-3216™) was purchased from ATCC. Mouse MSCs and hiMSCs were cultured in MesenCult™ Expansion Kit (Mouse) (05513, STEMCELL) and MesenCult™-ACF Plus Culture Kit (05448, STEMCELL), respectively, following the manufacturer's instructions and additionally supplemented with 1% penicillin-streptomycin (P/S). For hiMSCs, culture plates were coated with the animal component-free cell attachment substrate provided in the kit according to the manufacturer's instructions prior to cell seeding. HEK293T cells were cultured in Dulbecco's modified Eagle's medium (DMEM, Gibco, 2051526) supplemented with 10% fetal bovine serum (FBS) and 1% P/S. Cells were formed into spheroids prior to loading into the device.

#### Spheroid formation for encapsulation

On the day before encapsulation, cells were detached from culture plates using TrypLE (Gibco) for mouse MSCs and Animal Component-Free Cell Dissociation Kit (STEMCELL Technologies) for hiMSCs, according to the manufacturer's instructions at 37 °C. Then $2 \times 10^6$ cells in 3 mL of culture medium were seeded into each well of a 12-well suspension plate (CELLTREAT) and placed on an orbital shaker (Benchmark) at 100 rpm overnight to induce the spheroid formation.

#### Preparation of fibrin-collagen blended biodegradable matrix

A fibrin–collagen blended hydrogel was prepared as follows (per 1 mL): 319.7 µL of ultrapure water, 100 µL of 10× M199 (final 1×), and 25 µL of HEPES (final 25 mM) were combined. Collagen type I (316 µL; stock 9.48 mg $mL^{-1}$, final 3 mg $mL^{-1}$) was then added, and the pH was adjusted to approximately 7.4 by incremental addition of 1 N NaOH (~10 µL increments) until the solution turned deep orange. Subsequently, 100 µL of KnockOut™ serum replacement (final 10%), 100 µL of fibrinogen (stock 30 mg $mL^{-1}$, final 3 mg $mL^{-1}$), and 30 µL of fibronectin (stock 1 mg $mL^{-1}$, final 30 µg $mL^{-1}$) were added. The desired volume of matrix was first mixed with cells, and thrombin (100 U $mL^{-1}$) was then added to a final concentration of 6 U $mL^{-1}$ immediately before loading into the device.

#### Rapamycin preparation and injection

For rapamycin administration, a 50 mg $mL^{-1}$ rapamycin (J62473, Thermo Fisher Scientific) stock solution was prepared in 100% ethanol and stored at −80 °C. The stock solution was diluted to a final concentration of 1 mg $mL^{-1}$ in a vehicle composed of 5% PEG 400 (06855, Sigma) and 5% Tween 80 (BDH7781-2, BDH Chemicals) in deionized water. Mice received daily intraperitoneal injections of rapamycin at 3 mg $kg^{-1}$ body weight beginning on the day of transplantation.

**Synthesis and characterization of SB-diol and zwitterionic polyurethane**

3-(Butylbis(2-hydroxyethyl)ammonio)propane-1-sulfonate (SB-Diol) was synthesized as follows. N-butyldiethanolamine (30 g, 186 mmol), 1,3-propanesultone (25 g, 205 mmol), and dichloromethane (50 mL) were added to a 250 mL round-bottom flask. The mixture was stirred under an argon atmosphere for 48 h at 40 °C. The product was precipitated in diethyl ether and washed with diethyl ether three times. Afterward, the solvent was removed using a rotary evaporator to obtain white powder. Zwitterionic polyurethane (ZPU) was then synthesized as follows. PTMG was dried in a vacuum oven prior to use. SB-Diol (2.84 g, 10 mmol) and PTMG (10 g, 5 mmol) were dissolved in DMSO (200 mL) at 80 °C under an argon atmosphere. HDI (5.046 g, 30 mmol) was then added dropwise into the flask, followed by four drops of $Sn(Oct)_2$ catalyst. The mixture was stirred vigorously at 80 °C for 90 min. The mixture was then allowed to cool for 90 min, after which 1,4-diaminobutane (1.322 g, 15 mmol) was added dropwise and stirred for 24 h at 80 °C under an argon atmosphere. The molar ratio of (SB-Diol + PTMG):HDI:1,4-diaminobutanewas set as 1:2:1.A control polyurethane (PU) was synthesized under identical conditions without SB-Diol (molar ratio of PTMG:HDI:1,4-diaminobutanewas set as 1:2:1). The polymer solution was then precipitated in deionized (DI) water and then washed with DI water three times. The resulting white powder was further precipitated in diethyl ether and washed with diethyl ether three times. The solvent was removed using a rotary evaporator to obtain ZPU powder. The chemical structures of the SB-Diol monomer, control PU, and ZPU were confirmed by proton nuclear magnetic resonance. $^1H$ NMR ($D_2O$, 500 MHz, ppm): δ 3.97 (t, 4H), 3.54 (m, 6H), 3.4 (t, 2H), 2.91 (t, 2H), 2.15 (m, 2H), 1.68 (m, 2H), 1.34 (m, 2H), 0.90 (t, 3H) for SB-Diol monomer. $^1H$ NMR (DMSO-d6) for control PU and ZPU polymer.

**Characterization of electrospun zwitterionic polyurethane nanofibrous membranes**

The morphology of electrospun ZPU membranes was observed using a scanning electron microscope (Zeiss LEO 1550 (Keck) SEM). Fiber diameter was quantified using ImageJ software. Wall thickness was measured from cross-sections of the cylindrical membrane using an optical microscope (EVOS AMF4300). Fourier transform infrared (FT-IR) spectra of control PU and ZPU membranes were acquired using a Bruker Vertex V80V vacuum FT-IR spectrometer over a wavenumber range of 400–4,000 $cm^{-1}$ with 64 scans. For tensile testing, all samples were soaked in DI water prior to testing. The membrane was mounted on a dynamic mechanical analysis instrument (Instron, Model 5943) with a gauge length of ~1.5 cm between the clamps. Tensile testing was conducted at a rate of 1% $s^{-1}$ (1% original length per second) at room temperature. Stress (MPa) and strain (%) were calculated by the instrument software. Young's modulus was determined from the slope of the stress–strain curve in the linear elastic region between 10% and 20% strain. $n = 3$ samples were tested.

**Cell characterization**

To evaluate MSC surface marker expression before and after lentiviral engineering, flow cytometry was performed. Briefly, detached cells were washed with PBS, blocked with staining buffer (2% BSA in PBS), and incubated with the indicated antibodies (1:100 dilution) for 30 min at 4°C in the dark. After washing with PBS, cells were fixed with 2% paraformaldehyde for 10 min on ice, washed twice, and resuspended in staining buffer for acquisition on an Attune NxT flow cytometer (Thermo Fisher Scientific). The following antibodies were purchased from BioLegend: IgG isotype control (406001), anti-Sca-1 (108105), anti-CD29 (102205), anti-CD44 (103021), anti-CD31 (102405), anti-CD117 (105805), and anti-CD45 (103107). To assess the proliferative capacity of MSCs before and after lentiviral engineering and clonal selection, cumulative population doubling level (CPDL) analysis was performed over eight consecutive passages. At each passage, $0.5 \times 10^6$ cells were seeded in a T75 flask (Celltreat) and cultured for 72 h. Cells were then harvested and counted using a hemocytometer. The population doubling level (PDL) at each passage was calculated using the following equation:

$$PDL = 3.32 \times \log_{10}(N\_h / N\_s)$$

where N_h is the number of cells at harvest and N_s is the number of cells seeded. The CPDL was determined by cumulative addition of PDLs across successive passages.

**Anti-PGT121 antibody quantification**

Serum was collected as described. Anti-PGT121 antibody levels were measured by indirect ELISA following the same plate preparation, incubation, and detection protocol described for PGT121 quantification, with the following modifications: plates were coated with PGT121 at 1 μg $mL^{-1}$, serially diluted serum samples (1:50, 1:250, 1:1250, 1:6250, 1:31250, and 1:156250) were used as the analyte, serum from naive BALB/c mice served as a negative control, and bound anti-PGT121 antibodies were detected with HRP-conjugated goat anti-mouse IgG (H+L) (115-035-166, Jackson ImmunoResearch) diluted 1:5,000.

**Histological analysis**

Devices from *in vitro* experiments or those retrieved from *in vivo* experiments were fixed in 4% paraformaldehyde, dehydrated through a graded ethanol series and xylene, and embedded in paraffin. Paraffin blocks were sectioned at 5 µm using a microtome (Rankin Basics). For hematoxylin and eosin (H&E) staining, sections were deparaffinized by heating at 60 °C for 1 h followed by immersion in xylene, rehydrated through a graded ethanol series, and stained with hematoxylin (GHS216, Sigma-Aldrich) and eosin (HT110116, Sigma-Aldrich). Sections were then dehydrated through graded ethanol solutions, mounted with coverslips using mounting medium (17987-01, EMS). For immunofluorescence staining, sections were deparaffinized and rehydrated as described above. Antigen retrieval was performed by incubating sections in proteinase K (0.02 mg $mL^{-1}$ in 1× TE buffer; SRE0047, Sigma-Aldrich) at 37 °C for 20 min, followed by cooling at RT for 10 min. Sections were washed three times with PBST, permeabilized with 1% BSA and 0.25% Triton X-100 in PBS for 10 min, washed three times, and blocked with 5% BSA and 0.1% Triton X-100 in PBS for 1 h at RT. Sections were incubated with primary antibodies overnight at 4 °C, washed three times with PBST, and incubated with fluorophore-conjugated secondary antibodies for 1 h at RT in the dark. Nuclei were counterstained with 4′,6-diamidino-2-phenylindole (DAPI) for 5 min, and slides were mounted with coverslips using fluorescence mounting medium (S36936, Invitrogen). Images were acquired using a fluorescence microscope (APX100, Olympus). Rabbit anti-GFP (300-005-245, Jackson ImmunoResearch, 1:100) was used for primary antibody and Alexa Fluor 488-conjugated alpaca anti-rabbit IgG (H+L) (611-545-215, Jackson ImmunoResearch, 1:400) was used for secondary antibody.

**Cryopreservation and thawing of cell-loaded devices**

The experimental timeline is described in Fig. S6B. Cell-loaded devices were submerged in 2 mL of cryopreservation medium (12648010, Gibco) in a cryogenic vial (66021-942, VWR) and placed in a controlled-rate freezing container at −80 °C overnight. Vials were then transferred to liquid nitrogen for long-term storage. For thawing, cryogenic vials were incubated in a 37 °C water bath for approximately 1 min, and devices were washed with culture medium and returned to standard culture conditions.

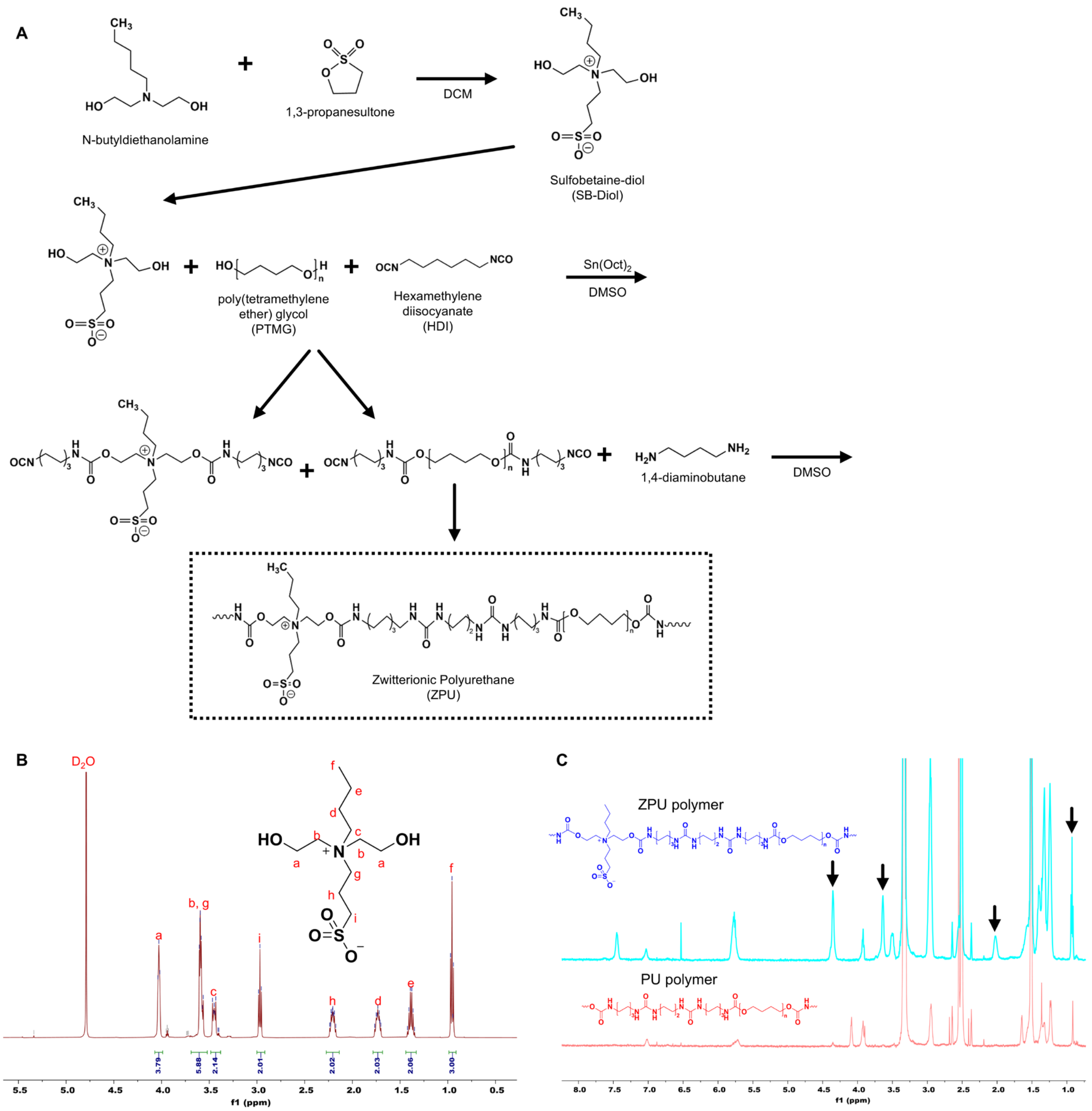


**Supplementary Fig. 1. (A)** Synthetic scheme for ZPU polymer, showing chemical structures of all precursors and intermediates. **(B)** $^{1}H$ NMR spectrum of SB-Diol monomer (500 MHz, $D_2O$). **(C)** $^{1}H$ NMR spectra of ZPU and PU polymers (500 MHz, DMSO-$d_6$). Arrows indicate peaks corresponding to sulfobetaine groups.

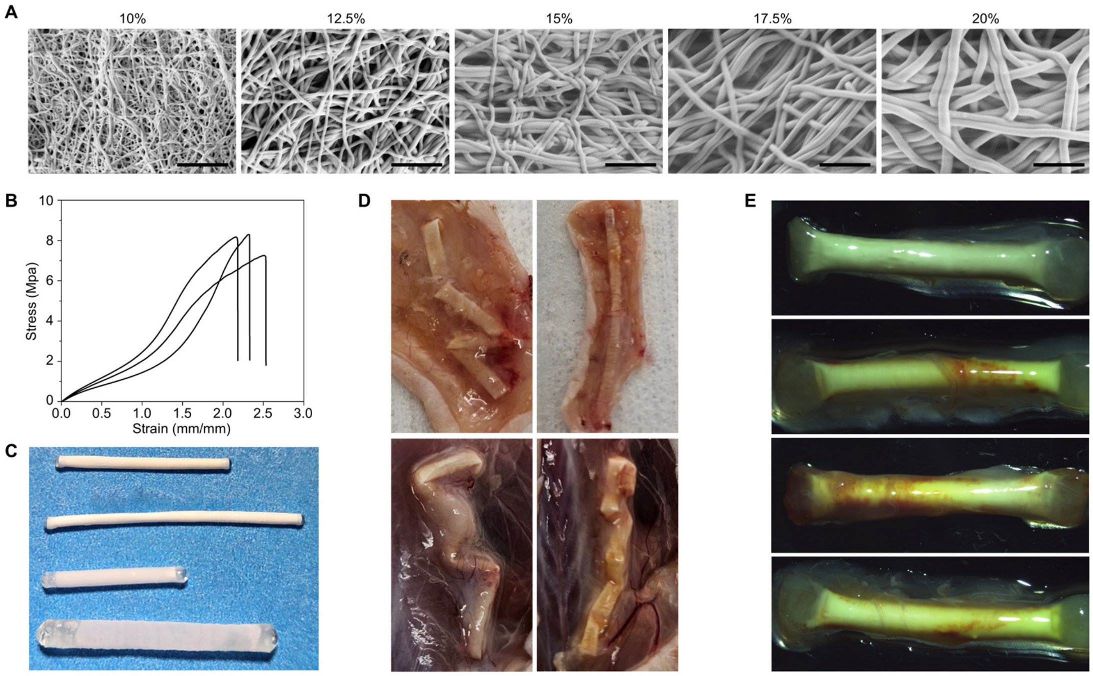


**Supplementary Fig. 2. (A)** SEM images of electrospun ZPU membranes fabricated from various ZPU solution concentrations (10, 12.5, 15, 17.5, and 20% w/v in HFIP). Scale bar, 5 µm. **(B)** Stress–strain curves of ZPU membranes fabricated from 10% (w/v) ZPU solution (*n = 3*). **(C)** Photographs of ZPU devices with varying diameters (1.2, 1.2, 1.6, and 2.8 mm) and lengths (2, 3, 1.5, and 2.5 cm), from top to bottom. **(D)** Additional photographs of retrieved Ver 1 devices (~75 µm wall, thermal sealing) at 8 weeks post-implantation in the dorsal subcutaneous space of BALB/c mice (*n = 5*). **(E)** Additional optical micrographs of retrieved Ver 2 devices (~156 µm wall, thermal sealing) at 4 weeks post-implantation in the dorsal subcutaneous space of BALB/c mice (*n = 5*).

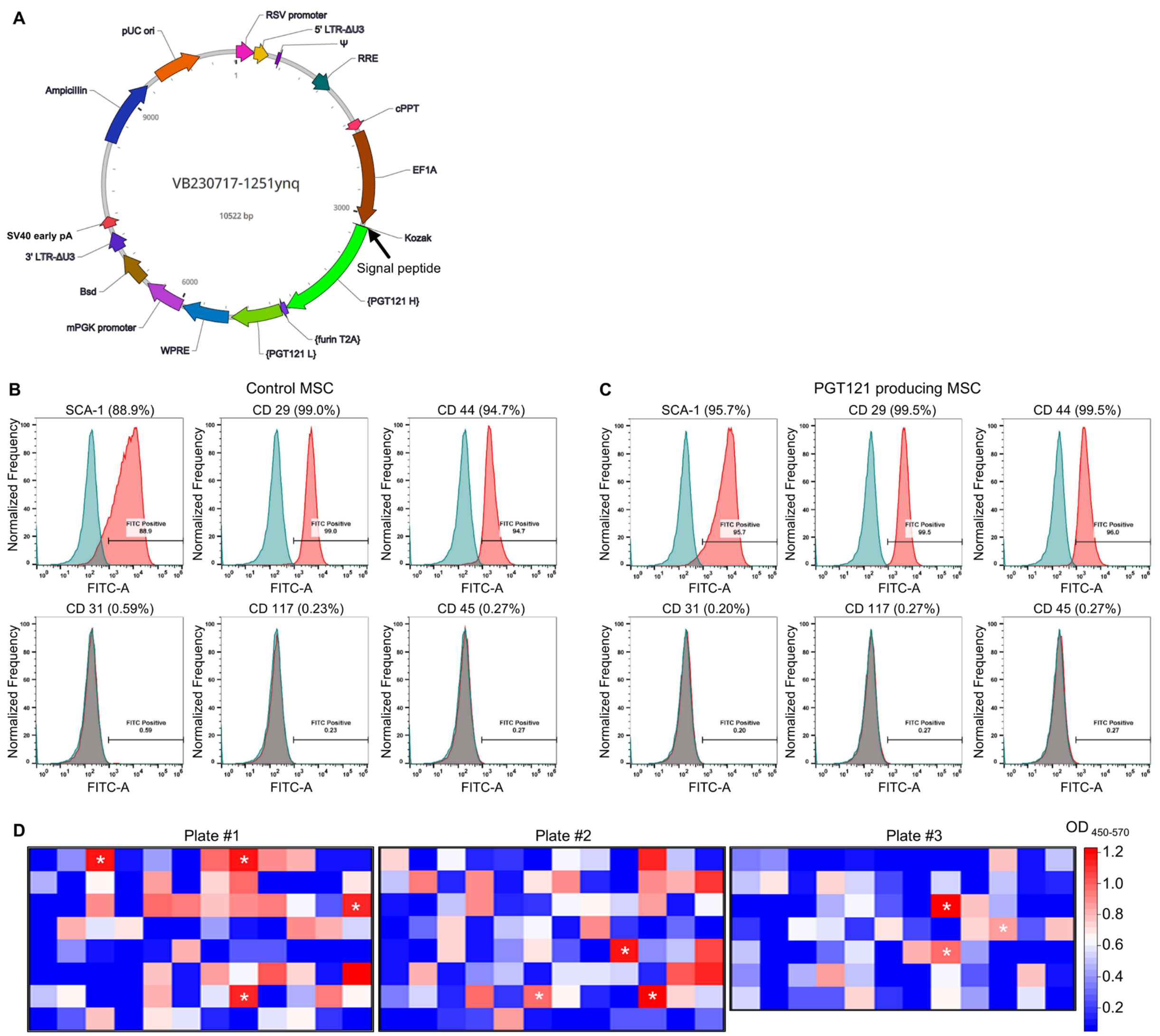


**Supplementary Fig. 3. (A)** Schematic of the lentiviral vector construct used for engineering cells to produce PGT121. **(B,C)** Flow cytometry histograms of **(B)** control MSCs and **(C)** PGT121-producing MSCs stained for MSC surface markers, including positive markers (Sca-1, CD29, CD44) and negative markers (CD31, CD117, CD45). Blue, isotype control; red, stained cells. **(D)** Heat map showing PGT121 production levels from 276 wells (three 96-well plates). Transduced MSCs were single-cell sorted into individual wells, expanded, and conditioned medium from each well was assayed by ELISA.

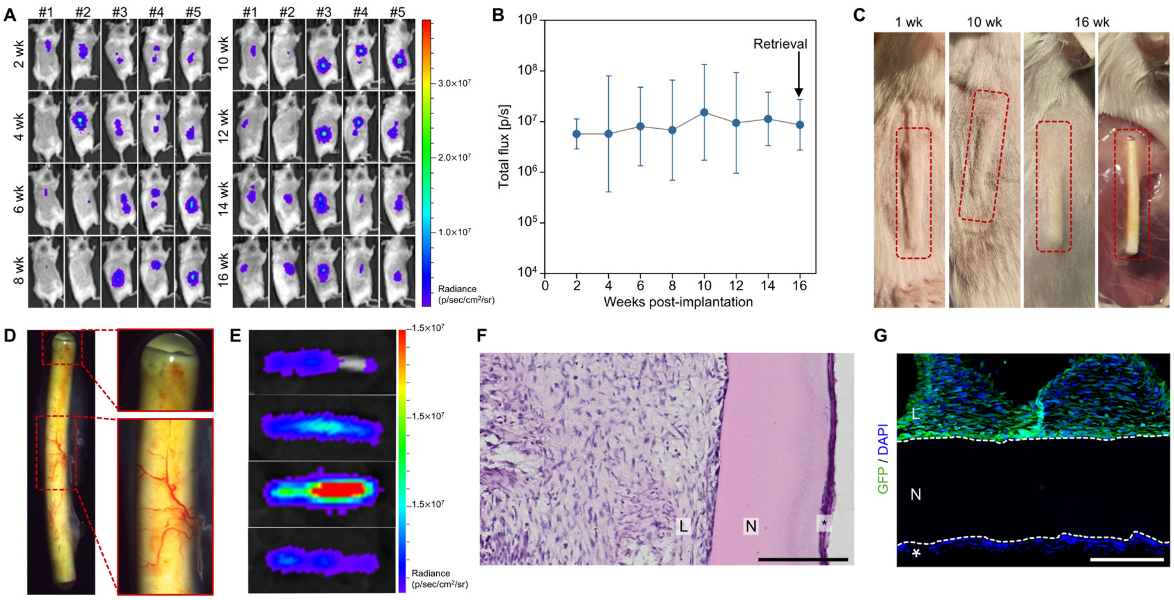


**Supplementary Fig. 4. (A–G)** Devices loaded with GFP/Luc-MSCs were implanted into the dorsal subcutaneous space of BALB/c mice (*n* = *5*). **(A)** Bioluminescence images of mice with implanted devices at 2, 4, 6, 8, 10, 12, 14, and 16 weeks post-implantation and **(B)** corresponding quantification. **(C)** Photographs of the implantation site (dorsum) at 1, 10, and 16 weeks post-implantation, and of exposed devices after skin incision at 16 weeks. **(D)** Optical micrographs of a device retrieved at 16 weeks. **(E)** Bioluminescence images of retrieved devices at 16 weeks post-implantation. **(F)** H&E-stained cross-sectional images of a device retrieved at 16 weeks. Scale bar, 200 µm. **(G)** Immunofluorescence images of cross-sectioned devices retrieved at 16 weeks. GFP (green) , DAPI (blue). Scale bar, 200 µm. In **(F)** and **(G)**: L, device lumen; N, nanofibrous membrane; *, fibrotic layer.

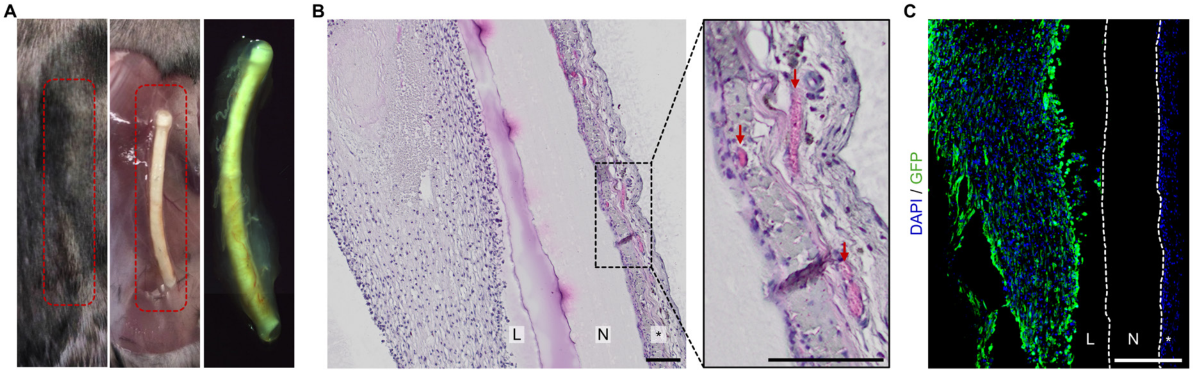


**Supplementary Fig. 5. (A–C)** Devices loaded with PGT121/GFP/Luc MSCs were implanted into the dorsal subcutaneous space of RAG2-KO mice (*n* = *8*). **(A)** Photographs of the implantation site (dorsum), exposed devices after skin incision, and optical micrograph of a retrieved device at 8 weeks post-implantation (left to right). **(B)** H&E-stained cross-sectional images of a device retrieved at 16 weeks. Scale bar, 100 µm. Arrows indicate vascular lumens. **(C)** Immunofluorescence images of devices retrieved at 16 weeks. GFP (green) and DAPI (blue). White dashed lines indicate the boundary of the nanofibrous membrane. In **(B)** and **(C)**: L, device lumen; N, nanofibrous membrane; *, fibrotic layer.

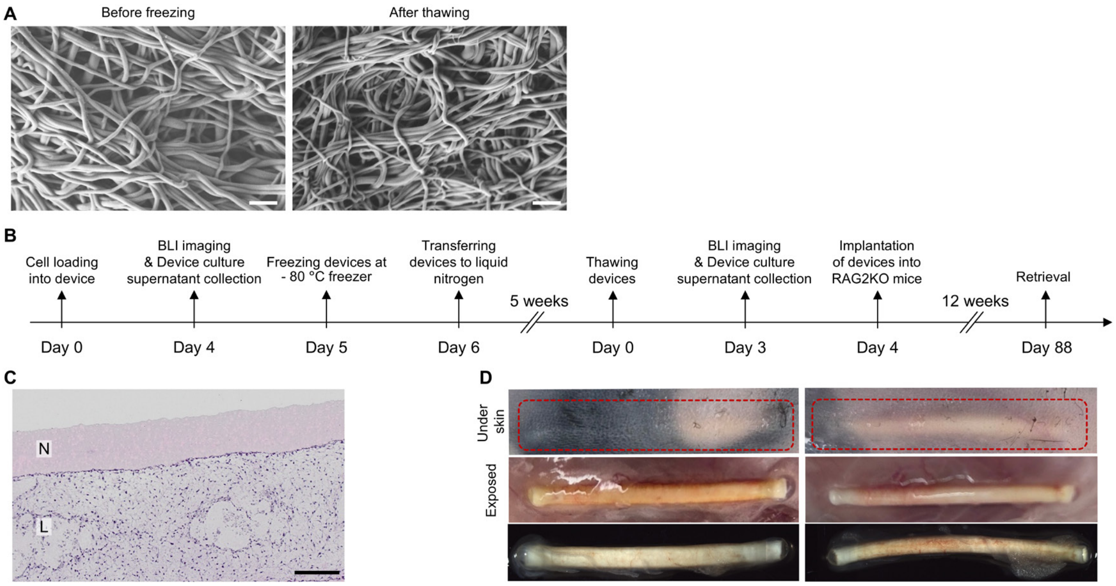


**Supplementary Fig. 6. (A)** SEM images of electrospun ZPU membranes before freezing and after storage in liquid nitrogen for 5 weeks followed by thawing. Scale bar, 2 µm. **(B)** Experimental timeline including bioluminescence imaging, *in vitro* productivity evaluation, cryopreservation and thawing, implantation, and retrieval for *ex vivo* evaluation. **(C)** H&E-stained cross-sectional images of devices loaded with PGT121/GFP/Luc-MSCs after a freeze-thaw cycle followed by *in vitro* culture. Scale bar, 200 µm. **(D)** Additional representative images at 12 weeks post-implantation. Photographs of the transplantation site (dorsum), exposed devices after skin incision, and optical micrographs of a retrieved device at 12 weeks post-transplantation (top to bottom) ($n = 3$).

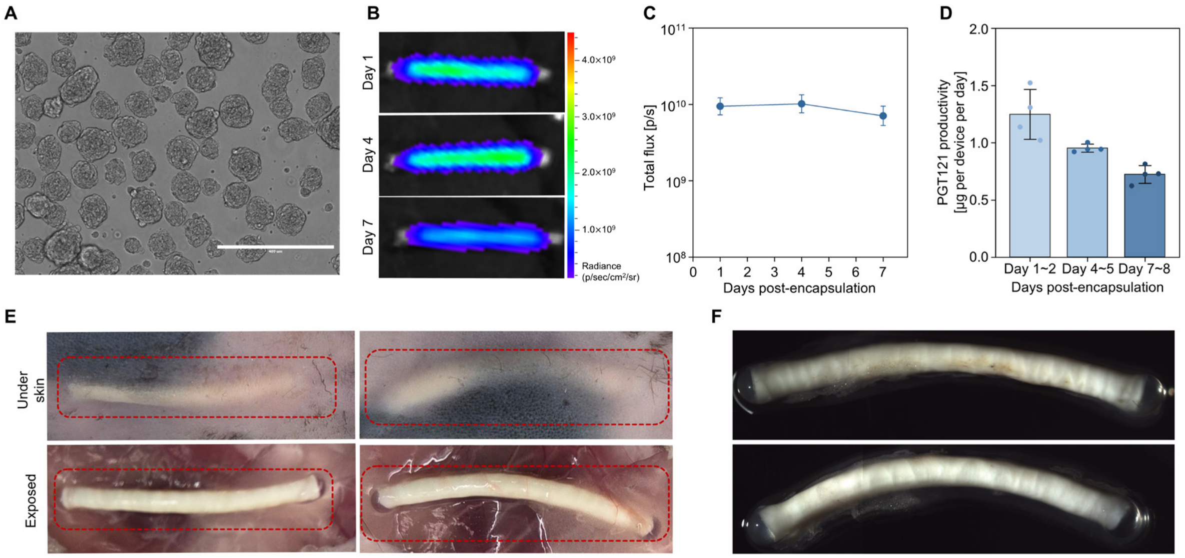


**Supplementary Fig. 7.** **(A)** Optical micrographs of hiMSC spheroids. Scale bar, 400 µm. **(B)** Bioluminescence images of devices loaded with 8 × $10^6$ PGT121/GFP/Luc-hiMSCs and **(C)** corresponding quantification ($n = 4$). **(D)** *In vitro* per-device PGT121 productivity of the same devices over 8 days of culture ($n = 4$). Conditioned medium was collected at days 1–2, 4–5, and 7–8 post-encapsulation. **(E, F)** Devices loaded with PGT121/GFP/Luc-hiMSCs were implanted into the dorsal subcutaneous space of RAG2-KO mice. **(E)** Additional photographs of the implantation site (top) and exposed devices after skin incision (bottom) at 12 weeks. **(F)** Additional optical micrographs of a retrieved device at 12 weeks.

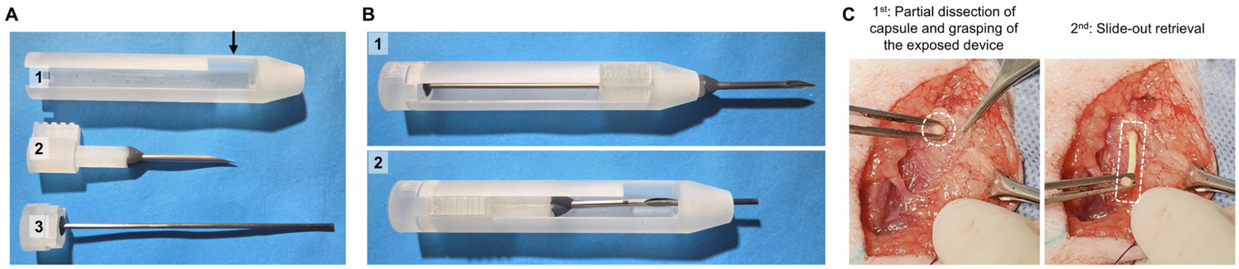


**Supplementary Fig. 8.** **(A)** Photographs of the scaled-up applicator for subcutaneous implantation in Göttingen minipigs, disassembled into three components: (1) body (arrow indicates locking notch), (2) needle with slider, and (3) obturator. **(B)** Photographs of the applicator: (1) slider locked, and (2) slider unlocked and needle retracted to deploy the device. **(C)** Retrieval procedure. Sequential steps (left to right): skin incision, partial dissection of the fibrotic layer, grasping of the exposed device tip, and slide-out retrieval.
